# IKAROS facilitates antigen escape in the face of CD19- and CD22-targeted therapies for B-cell acute lymphoblastic leukemia

**DOI:** 10.1101/2024.11.01.621347

**Authors:** Pablo Domizi, Jolanda Sarno, Astraea Jager, Milton Merchant, Kaithlen Zen B Pacheco, Sean A. Yamada-Hunter, Maria Caterina Rotiroti, Yuxuan Liu, Reema Baskar, Warren D. Reynolds, Brian J. Sworder, Bita Sahaf, Sean C. Bendall, Charles G. Mullighan, Ash A. Alizadeh, Allison B. Leahy, Regina M. Myers, Bonnie Yates, Hao-Wei Wang, Nirali N. Shah, Robbie G. Majzner, Crystal L. Mackall, Stephan A. Grupp, David M. Barrett, Elena Sotillo, Kara L. Davis

## Abstract

Relapse due to antigen escape is a major cause of treatment failure for patients with B-cell malignancies following targeted immunotherapies, including CD19- and CD22-directed chimeric antigen receptor T (CAR T) cells. To identify tumor intrinsic factors associated with antigen loss, we performed single-cell analyses on 61 primary patient samples or patient-derived xenografts from patients with B-cell acute lymphoblastic leukemia (B-ALL) treated with CAR T cells. We identified that low levels of the transcription factor IKAROS in pro-B-like B-ALL cells before CAR T treatment are associated with antigen escape. We demonstrate that IKAROS^low^ B-ALL cells lose features of B cell identity and resemble progenitor cells based on their epigenetic and transcriptional state, resulting in the downregulation of B-cell immunotherapy antigens, including surface expression of CD19 and CD22. We find that modulation of CD19 and CD22 protein expression is IKAROS dose-dependent and reversible. Further, we demonstrate that IKAROS^low^ cells are resistant to CD19- and CD22-targeted therapies. Together, we describe a novel role for IKAROS in the regulation of B-cell immunotherapy targets and the risk of antigen escape relapse, identifying it as a potential prognostic target.

**Highlights:**

- IKAROS^low^ pro-B-like B-ALL cells are associated with CD19^neg^ relapse
- IKAROS^low^ B-ALL cells resemble progenitor cells and have lower B-cell commitment
- IKAROS modulates CD19 and CD22 surface expression in a dose-dependent and reversible manner
- IKAROS^low^ B-ALL cells are more resistant to CD19- and CD22-targeted therapies

**Graphical abstract:** 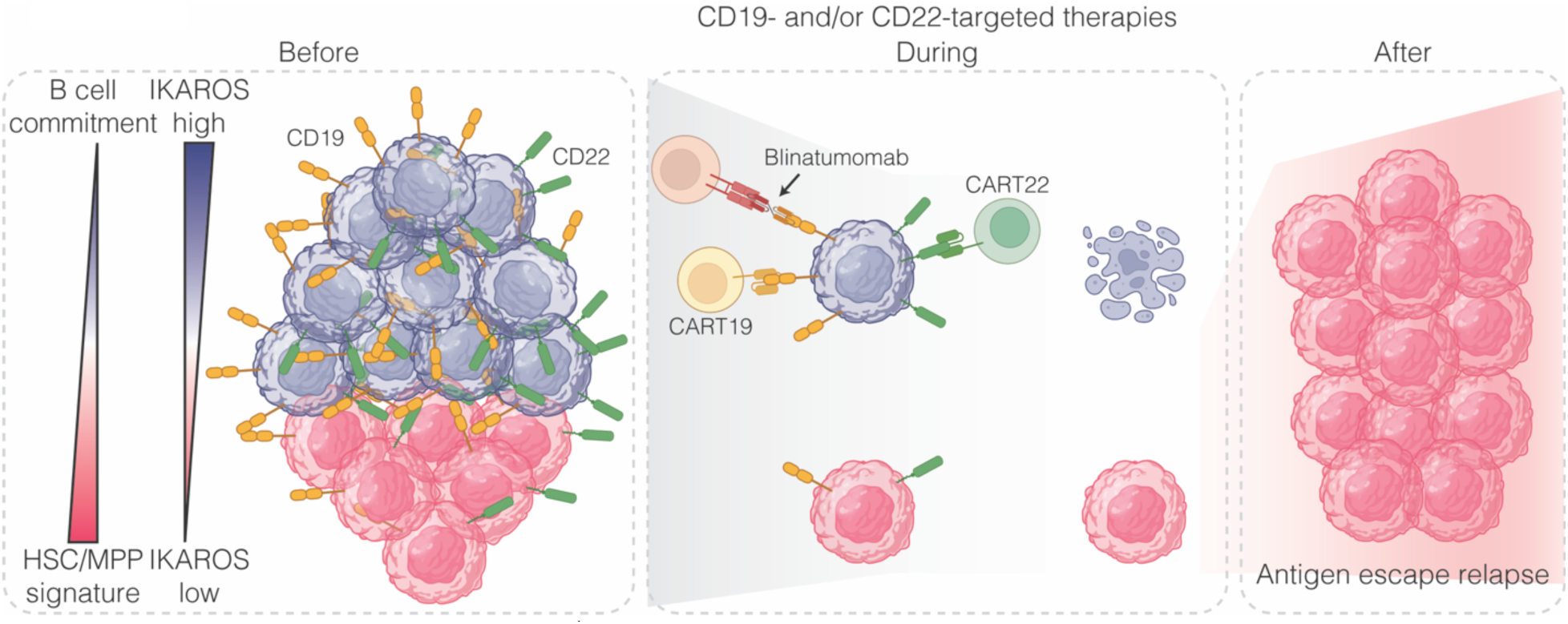

Before immunotherapy, IKAROS^low^ pro-B-like B-ALL cells possess chromatin and gene expression states poised for loss of B-cell identity while maintaining expression of CD19 and CD22. Under immune pressure, IKAROS^high^ cells maintain their antigen expression, making them more susceptible to T cell-mediated killing. Conversely, IKAROS^low^ cells are more likely to downregulate their antigen expression, giving them a relative advantage to escape immunotherapies, resulting in antigen escape relapse.

## Introduction

B-cell acute lymphoblastic leukemia (B-ALL) is the most common childhood cancer, accounting for 25% of cancer diagnoses in children and adolescents up to 19 years of age^1^. Approximately 15% of children and young adults diagnosed with ALL will relapse, and 50% of children who relapse will die, making relapsed B-ALL the second leading cause of cancer-related death for children in the U.S^1^. Adult B-ALL has a dismal prognosis, with only 40% of patients alive 5 years from diagnosis^2^. CD19-directed CAR T (CART19) cells are now standard therapy for children and adults with relapsed or refractory B-ALL, achieving remission rates of 70 – 90%^3,4^. Unfortunately, around 50% of initial responders will eventually relapse, many with CD19 loss^5,6^. Similarly, CD19^neg^ relapses have been reported in patients treated with blinatumomab, an anti-CD3/CD19 bispecific T cell engager^7,8^. For patients suffering CD19^neg^ relapse, CD22-targeted therapies, either inotuzumab ozogomycin or CD22-directed CAR T (CART22) cells, have emerged as a salvage option^9^. However, CD22 downregulation has limited durable remissions^10^. Several mechanisms of CD19 loss have been reported, including truncated CD19 mutations^11^, disruption of CD19 trafficking to the cell surface^12,13^, *CD19* mRNA mis-splicing^14,15^, and lineage switch^16^. Fewer studies have focused on the mechanisms behind CD22 downregulation, but transcriptional downregulation and alternative splicing have also been reported^17^. In several studies, CD19 loss was accompanied by CD22 downregulation^18–22^. Recovery of CD19 and CD22 expression has been reported after relief of immune pressure ^17,18,23,24^.

To discover cell-intrinsic factors associated with antigen loss, we analyzed 39 pre- and post-CART19 patient-derived xenografts (PDXs) from 25 patients using mass cytometry (CyTOF) and single-cell RNA and antibody tag sequencing (CITE-seq). In pre-CART19 samples, we identified CD19^pos^ pro-B-like tumor cells with low expression of the transcription factor IKAROS to be associated with CD19^neg^ relapse. We determined that IKAROS regulates CD19 expression in B-ALL, large B cell lymphoma (LBCL), and chronic lymphoblastic lymphoma (CLL) models. IKAROS^low^ cells demonstrate a wholesale change in chromatin and transcriptional state, shifting their identity away from B cells and moving towards myeloid and progenitor states. This loss of B cell identity manifests with decreased CD19 and CD22 surface expression. Consistent with this, we confirmed low IKAROS levels are also associated with CD22^low^ relapse in an independent cohort of 11 r/r B-ALL patients treated with CART22. IKAROS-mediated decrease in CD19 and CD22 surface expression confers a survival advantage for IKAROS^low^ B-ALL cells against CD19- and CD22-targeted therapies. Together, we describe a novel role for IKAROS in immunotherapy target regulation and implicate pre-existing IKAROS^low^ cells at risk of antigen loss under the pressure of CD19- or CD22-targeted therapies in B-ALL patients.

## Results

### CD19 loss is accompanied by loss of B cell features

To identify tumor-intrinsic factors associated with CD19^neg^ relapse following CAR T therapy, we profiled 35 patient-derived xenograft (PDX) samples from pediatric and adult leukemia patients treated with 19.BBz CAR T cells using CyTOF (Figure 1A). This cohort includes pre-CART19 PDX samples from patients who achieved durable complete remission (CR; n = 6), underwent CD19^neg^ (n = 11) or CD19^pos^ (n = 4) relapse, or were non-responders (n = 4). We also analyzed paired post-CART19 relapse PDX samples from 14 of these patients, including those with CD19^neg^ relapse (n = 8), CD19^pos^ relapse (n = 2), or non-response (n = 4). Evaluation of CD19 expression in pre-CART19 PDX samples did not show differences across the clinical groups (Figure 1B).

**Figure 1:**
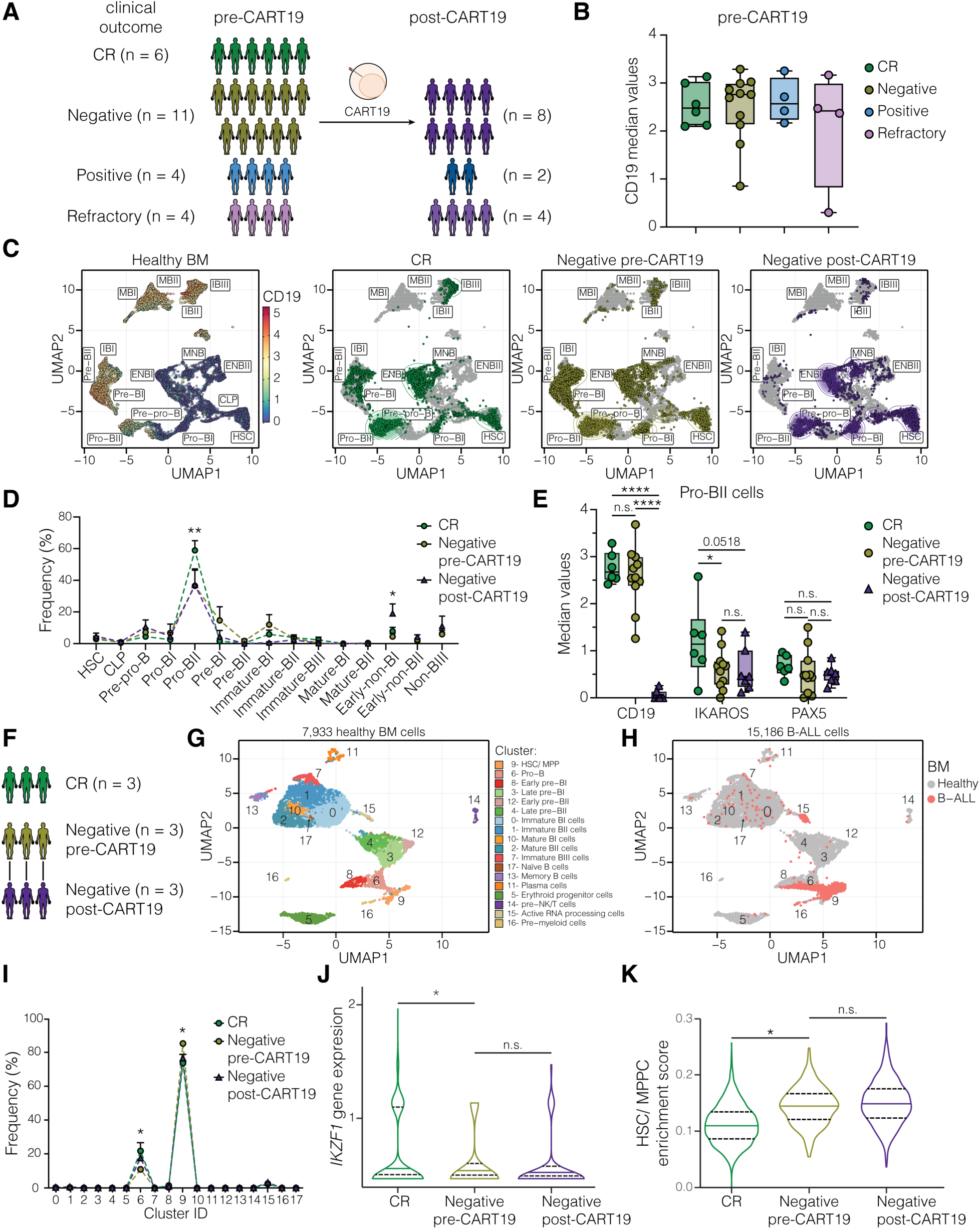
CD19 loss is associated with loss of B cell identity. **(A)** CART19 cohort of PDX samples analyzed by CyTOF. **(B)** Median CD19 expression in pre-CART19 samples. **(C)** UMAP based on developmental classifier protein expressions in Lin-/ B+ fraction of healthy BM (left) and projection of tumor cells from pre-CART19 patients that achieved durable CR, suffered CD19^neg^ relapse, and post-CD19 loss, respectively. IBI = Immature-BI, IBII = Immature-BII, IBIII = Immature-BIII, MBI = Mature-BI, MBII = Mature-BII, ENBI = Early-non-BI, ENBII = Early-non-BII, and NBIII = Non-BIII. **(D)** Developmental classification of samples in (C). The pro-BII population is significantly more abundant in the CR group, while early-non-BI is significantly more abundant between the pre- and post-CD19^neg^ relapse groups. **(E)** Median protein expression of CD19, IKAROS, and PAX5 in pro-BII-like tumor cells. **(F)** Cohort of pre-and post-CART19 B-ALL samples analyzed by single-cell CITE-seq. **(G)** UMAP based on most variables genes in Lin-/ B+ fraction of healthy BM. Space occupied by healthy clusters is depicted. **(H)** Projection of B-ALL cells from 9 samples onto healthy BM-defined UMAP space. **(I)** Association of B-ALL cells to their closest healthy cluster, based on k-nearest neighbor assignment. (**J - K**) *IKZF1* gene expression (J) and single-cell enrichment score for HSC/MPP gene signature **(K)** in pro-B-like B-ALL cells (cluster 6). Boxplots in (B) and (E) show median with range from minimum to maximum values. Curves in (D) and (I) show mean ± SEM. Violin plots in (J) and (K) show median (solid line) and 25^th^ and 75^th^ quantile (dash lines). The statistical tests used were one-way ANOVA followed by Tukey’s multiple comparisons tests (B); two-way ANOVA followed by Tukey’s multiple comparisons tests (D), (E), and (I); and Wilcoxon rank sum test followed by Bonferroni’s multiple comparisons test in (J) and (K). *P<0.05, **P<0.01, ****P< 0.0001; not significant (n.s.), P>0.05.

We applied our B-cell developmental classifier to compare cell types across patients^25^. Consistent with our prior findings in de novo B-ALL^25^, most leukemic cells were classified within the pre-pro-B to pre-BI transitional populations, particularly as pro-BII-like B-ALL cells (Figures 1C, 1D). Interestingly, after CD19 loss, a significant fraction of B-ALL cells were classified as early-non-BI (Figure 1D). Early-non-BI cells are progenitor cells that do not express CD19 or other pan B-cell markers (Figure S1B). Enrichment in the early-non-BI population after CART19 was not observed in patients with CD19^pos^ relapse (n = 2) or refractory patients (n = 4; Figures S2A, S2B). This enrichment was not observed when comparing isogenic CD19 wild-type (WT) or knockout (KO) B-ALL cell lines (Figure S2C), suggesting that the sole loss of CD19 expression does not alter the developmental profile of B-ALL cells. Instead, in patients, CD19 loss is accompanied by the loss of additional B-cell features.

### Low levels of IKAROS in pro-B-like tumor cells are associated with CD19^neg^ relapse

Since pro-BII-like B-ALL cells were the most abundant across all patient groups (Figure 1D), we compared protein expression in pro-BII-like cells from patients who achieved durable CR or underwent CD19^neg^ relapse. While CD19 and PAX5 expression did not differ between pre-treatment groups, pro-BII-like cells from patients who would experience CD19^neg^ relapse had lower expression of the B-lineage transcription factor IKAROS (Figure 1E). This difference in IKAROS level was specific to pro-BII like cells. It was not observed in the bulk leukemia cells or any other population (Figure S2D).

To further explore the differences between patients who achieved durable CR and those who underwent CD19^neg^ relapse, we performed CITE-seq on cells from healthy BM (n = 1), pre-CART19 samples from patients who achieved CR (n = 3), and paired pre- and post-CD19^neg^ relapse samples (n = 3) (Figure 1F). We defined 18 cell populations in the healthy BM based on gene and protein expression profiles (Figures 1G and S3A). Then, we projected each B-ALL cell onto the healthy BM space (Figure 1H) and assigned its closest healthy population. Most B-ALL cells were associated with healthy hematopoietic multipotent progenitor cells (HSC/MPP, cluster 9) or pro-B cells (cluster 6) (Figure 1I). Although bulk samples and HSC/MPP-like B-ALL cells showed no differences in gene expression across clinical outcomes (Figures S3B and S3C), pre-CART19 pro-B-like B-ALL cells from patients with durable CR or CD19^neg^ relapse exhibited distinct gene signatures (Figure S3D). In particular, *IKZF1* expression was significantly lower in pre-CART19 CD19^neg^ relapse pro-B-like B-ALL cells (Figure 1J). This was not true in bulk samples or HSC/MPP-like cells (Figure S3E). There were no differences in *CD19* mRNA, CD19 protein, or *PAX5* mRNA levels in pre-CART19 pro-B-like B-ALL cells (Figure S3F), consistent with our findings at the protein level (Figure 1E). Pre-CART19 *IKZF1*^low^ pro-B-like B-ALL cells from patients who suffered CD19^neg^ relapse were enriched for an HSC/MPP gene expression signature, indicating these cells had less B-cell identity before CART19 administration (Figure 1K). These features, low levels of *IKZF1* expression and enrichment in HSC/MPP gene expression signature, were conserved after CD19 loss (Figures 1J and 1K).

### IKAROS is required to sustain B-cell identity and CD22 surface expression

To investigate the impact of IKAROS on CD19 surface expression, we targeted IKAROS in B-ALL cell lines using two separate short hairpin RNA (shRNA) sequences (KD1 and KD2, Figure 2A), or lenalidomide, a cereblon inhibitor that targets IKAROS for degradation^26^ (Figure 2B), or a combination of both methods (KD1 with or without 10 µM lenalidomide, Figure 2C). These experiments confirmed that decreased IKAROS levels reduced CD19 surface expression. Furthermore, we confirmed the relationship between IKAROS and CD19 surface expression in LBCL (Figures S4A, S4B, and S4C) and CLL (Figure S4D) models, suggesting that IKAROS modulation of CD19 surface expression is consistent across B cell malignancies.

**Figure 2:**
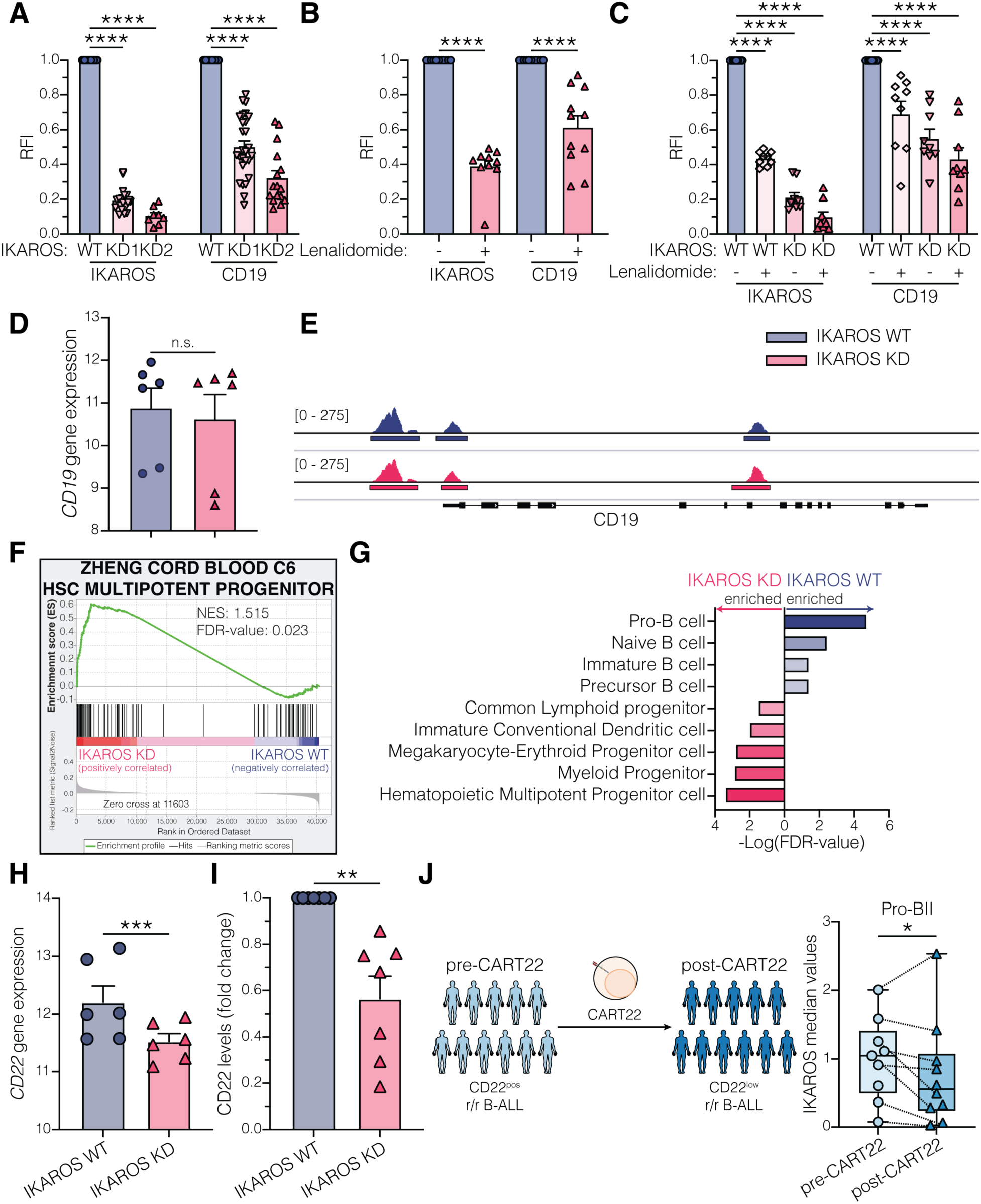
IKAROS regulates CD19 and CD22 surface expression. **(A - C)** Relative IKAROS and CD19 median levels in B-ALL cell lines (697, NALM6, NALM16, NALM20, REH, RS4;11, SUP-B15) transduced with lentivirus expressing scrambled or short hairpin RNA (shRNA) against *IKZF1* (A; n = 8 - 27), treated with DMSO or 10 µM lenalidomide (B; n = 11), or combining shRNA knockdown with or without lenalidomide treatment (C, n = 9). Proteins were measured by flow cytometry and normalized to scrambled transduced (A), DMSO-treated (B), or scrambled transduced and DMSO-treated (C) cells. RFI = relative fluorescence intensity. **(D)** *CD19* variance-stabilized transformed (vst) counts in IKAROS WT or KD B-ALL cells. **(E)** Accessibility profile of *CD19* promoter and gene from one representative IKAROS WT and KD B-ALL cell line pair. Other cell lines and their biological replicates can be found in Figure S5A. **(F)** GSEA results for Zheng Cord Blood C6 HSC/MPP gene signature^68^ in IKAROS WT and KD B-ALL cells. **(G)** Cell type enrichment analysis of genes with differentially accessible promotors in IKAROS WT or IKAROS KD B-ALL cells. **(H)** *CD22* vst counts in IKAROS WT or KD B-ALL cells. **(I)** Relative CD22 median levels in isogenic IKAROS WT or KD B-ALL cells (n = 7). Values were measured by flow cytometry or CyTOF and normalized to IKAROS WT condition. **(J)** Paired pre-CART22 and post-CD22^low^ relapse cohort of leukemic patient samples analyzed by CyTOF. IKAROS median levels in pro-BII-like tumor cells. RNA-seq and ATAC-seq experiments (D - H) were performed in 3 cell lines (NALM6, REH, and SUP-B15) with two biological replicates per cell line. Bar plots in (A - D) and (H - I) show mean ± SEM. Boxplots in (J) show median with range from minimum to maximum values. Statistical tests used were two-way ANOVA followed by Šidák’s multiple comparisons test (A - C); differential gene expression analysis based on the negative binomial distribution (D) and (H); paired t-test (I) and (J). *P<0.05, **P<0.01, ***P< 0.001, ****P< 0.0001; not significant (n.s.), P>0.05.

To understand how IKAROS modulates CD19 surface expression, we analyzed isogenic IKAROS wild-type (WT) and knockdown (KD) B-ALL cell lines using ATAC-seq and RNA-seq (Figure S4E). Transcriptomic data showed similar *CD19* mRNA levels in IKAROS WT and KD cells (Figure 2D). Chromatin accessibility at the *CD19* promoter and gene locus also showed no changes (Figures 2E and S5A). ATAC-seq analysis identified 1,250 and 4,037 peaks significantly more accessible in IKAROS WT and KD cells, respectively, consistent with IKAROS’ role as a transcriptional repressor^27^. The transcription factor binding sites in these differentially accessible peaks differed between IKAROS WT and KD cells (Figure S6A), indicating altered gene expression programs in IKAROS KD cells. Transcriptomic analysis confirmed differential expression of several transcription factors between IKAROS WT and KD cells (Figure S6B). IKAROS KD cells showed 168 genes with both more accessible promoters and higher expression, forming a network characterized by non-B lineage genes (red nodes in Figure S6C), consistent with a loss of B cell identity. Indeed, while the gene expression profile and chromatin landscape of IKAROS WT cells are consistent with pro-B cell identity (Figures 2G and S4G), IKAROS KD cells acquired a profile more characteristic of HSC/MPP cells (Figures 2F and 2G).

Given the re-wiring of transcriptional networks in IKAROS KD cells, we investigated the expression of other B-cell proteins. IKAROS KD cells showed reduced *CD22* mRNA and CD22 surface protein expression (Figures 2H and 2I). Clinical trials have targeted CD22 in patients who relapse after CD19-targeted therapies^10^. We analyzed 22 paired patient samples collected before CART22 administration or after CD22^low^ relapse to assess the clinical implications of IKAROS and CD22 interaction. Lower IKAROS levels were found in pro-BII-like B-ALL cells following CD22^low^ relapse (Figure 2J). These results suggest that IKAROS modulates CD19 and CD22 surface expression, and patients with IKAROS^low^ tumor cells might be at higher risk for antigen-loss relapse following both CD19- and CD22-targeted therapy.

### IKAROS regulates CD19 and CD22 surface expression in a dose-dependent and reversible manner

To fine-tune IKAROS levels without the toxicities associated with shRNA infection and lenalidomide treatment, we generated IKAROS-regulatable models by overexpressing a codon-optimized version of IKAROS fused to a degron tag^28^ and knocked out the endogenous *IKZF1* gene. In this system, in the absence of asunaprevir, a hepatitis C virus nonstructural protein 3 protease inhibitor, cells express high levels of IKAROS as the degron tag is removed. However, in the presence of asunaprevir, IKAROS is rapidly targeted for degradation. We generated seven models (IKAROS-degron KO1 – KO7) where IKAROS levels can be titrated using different concentrations of asunaprevir (Figure 3A). CD19 and CD22 surface levels decreased in an IKAROS dose-dependent manner (Figures 3B and 3C). We selected IKAROS-degron KO1 and KO2 models for further studies because they express the highest and lowest baseline CD19 surface expression, respectively. Upon asunaprevir withdrawal, IKAROS, CD19, and CD22 levels were restored (Figures 3D, 3E, and 3F), demonstrating reversible regulation of CD19 and CD22 surface expression by IKAROS.

**Figure 3:**
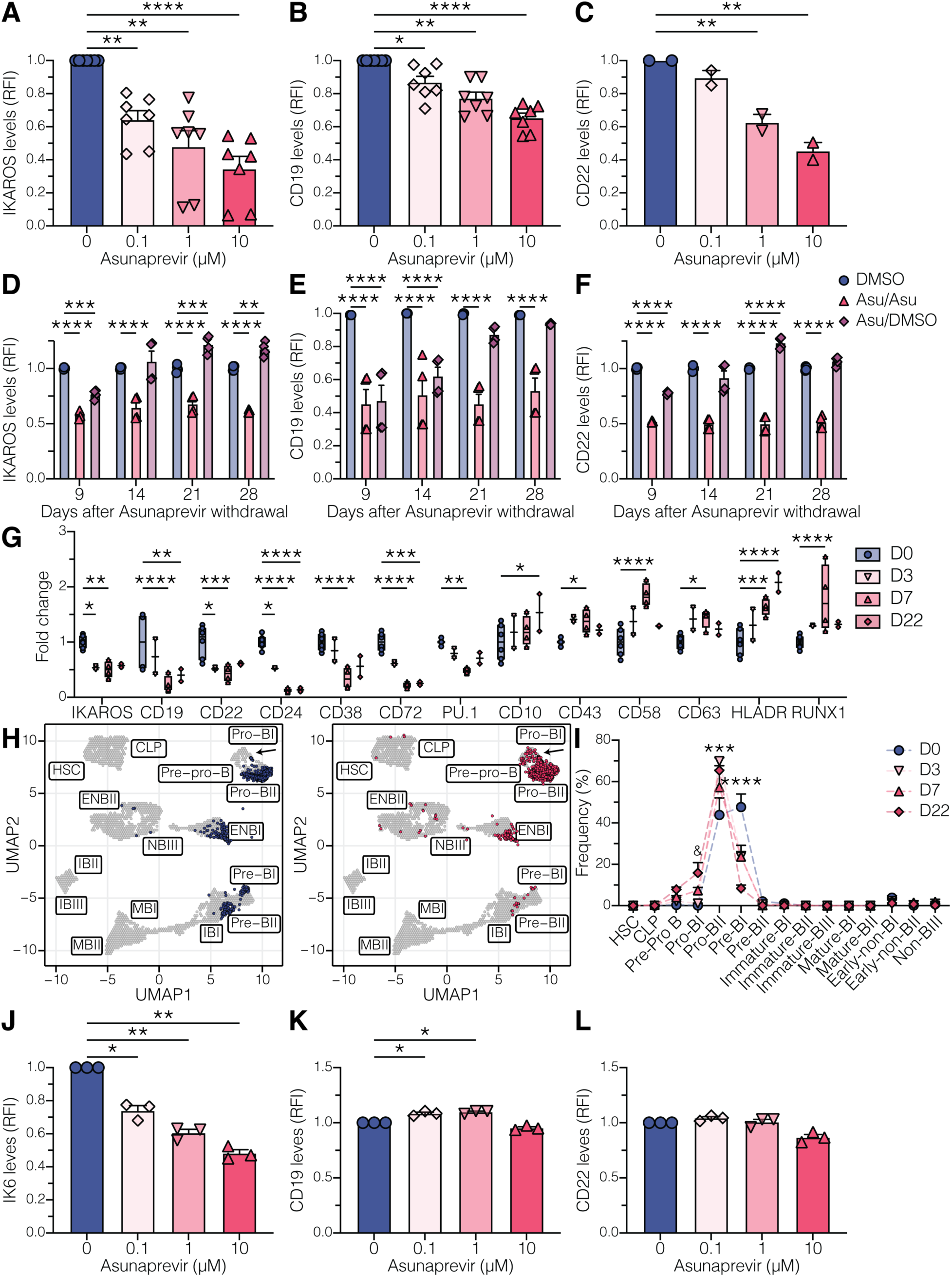
IKAROS regulates CD19 and CD22 surface expression in a dose-dependent and reversible manner. **(A - C)** Relative IKAROS (A, n = 7), CD19 (B, n = 7), and CD22 (C, n = 2) median levels in IKAROS-degron models treated with increasing doses of asunaprevir. Values were measured by flow cytometry and normalized to DMSO-treated (0 µM) condition. Dots represent the mean value from two technical replicates. RFI = relative fluorescence intensity. **(D - F)** IKAROS-degron models were treated with 10 µM asunaprevir for 7 days. Afterward, asunaprevir was withdrawn, and cells were either treated with DMSO (ASU/DMSO) or maintained with 10 µM asunaprevir (ASU/ASU) for an additional 28 days. As a control, IKAROS-degron models were treated with DMSO for 35 days. Relative levels of IKAROS (A), CD19 (B), and CD22 (C) were measured at 9, 14, 21, and 28 days post-asunaprevir withdrawal by flow cytometry and normalized to DMSO-treated condition. The experiment was performed in duplicate. RFI = relative fluorescence intensity. **(G)** IKAROS-degron models were treated with DMSO (D0; n = 6) or 10 µM asunaprevir for 3 (n = 2), 7 (n = 4), and 22 (n = 2) days. The protein profile was measured by flow cytometry and CyTOF and normalized to the D0 condition. **(H)** Projection of IKAROS-degron models treated with DMSO (left) or 10 µM asunaprevir (right) for 7 days onto the UMAP representation of healthy B cell developmental populations based on expression of developmental classifier protein expressions. **(I)** Developmental classification of samples in (G). “&” denotes p-value < 0.01 in Pro-BI population frequency between D0 and D22 conditions. The frequencies of Pro-BII and Pre-BI populations were statistically different (p-value < 0.001) in the D0 condition against D3, D7, and D22 conditions. **(J - L)** Relative IK6 (A, n = 3), CD19 (B, n = 3), and CD22 (C, n = 3) median levels in IK6-degron models treated with increasing doses of asunaprevir. Values were measured by flow cytometry and normalized to DMSO-treated (0 µM) condition. Dots represent the mean value from two technical replicates. RFI = relative fluorescence intensity. Bar plots in (A - F) and (J - L) show mean ± SEM. Boxplots in (G) show the median, with a range from minimum to maximum values. Curves in (I) show mean ± SEM. Statistical tests used were paired one-way ANOVA followed by Dunnett’s multiple comparisons test (A - C) and (J - L); and two-way ANOVA followed by Dunnett’s multiple comparisons test (D - G) and (I). *P<0.05, **P<0.01, ***P< 0.001, ****P< 0.0001.

CyTOF analysis of IKAROS-degron models treated with asunaprevir for 3, 7, or 22 days showed downregulation of CD19 and CD22 surface expression along with mature B-cell (CD24, CD38, CD72) and B-cell differentiation (PU.1) proteins in IKAROS^low^ conditions. Conversely, proteins in progenitor (CD10, CD43), myeloid (CD58, CD63, RUNX1), and antigen-presenting (HLA-DR) cells were upregulated (Figure 3G). These phenotypic changes promote a shift of IKAROS^low^ cells toward more progenitor and immature B-cell states (Figures 3H and 3I), confirming previous observations of the transcriptional and chromatin landscapes.

IKAROS alterations are associated with relapse in de novo B-ALL^29^. The most common IKAROS alteration is the deletion of the four DNA-binding domains, resulting in the dominant negative IK6 isoform^30^. To assess whether patients expressing the IK6 isoform—who are more likely to experience chemotherapy failure and subsequently receive CD19- or CD22-targeted therapies— are at higher risk for antigen escape relapse, we generated IK6-degron models (Figure S7A). Surface CD19 and CD22 molecules were similar among IKAROS-degron and IK6-degron models (Figure S7B). Interestingly, decreasing IK6 expression with asunaprevir treatment did not modify CD19 or CD22 surface levels in IK6-degron models (Figures 3J, 3K, and 3L). CyTOF protein profiling revealed that asunaprevir treatment did not alter the expression of CD38, CD58, CD72, or RUNX1, while the effects on CD24, PU.1, and HLA-DR expression were attenuated (Figure S7C). CD45 was the only protein that showed a distinct response to asunaprevir treatment, being upregulated in IK6-degron cells (Figure S7C). Similarly, IK6-degron models did not exhibit a developmental shift following asunaprevir treatment (Figure S7D). Consistent with the results of our model, in primary patient data, there are no differences in *CD19* or *CD22* gene or protein expression between patients with *IKZF1* deletions and those with other B-ALL subtypes (Figures S7E and S7F). These results suggest that patients with *IKZF1* deletions are not more susceptible to CD19 or CD22 downregulation, and in the context of CD19- and CD22-targeted therapies, the response is instead related to wild-type IKAROS dose and requires the DNA binding domains.

### Low levels of IKAROS confer resistance to CD19- and CD22-targeted therapies

Since antigen density is crucial for CAR T cell efficacy^31^, we investigated whether IKAROS^low^ cells have an advantage against CD19- and CD22-targeted therapies. First, we confirmed that low IKAROS levels significantly reduced the number of CD19 and CD22 molecules on the B-ALL cell surface (Figures 4A, 4B, and 4C). Then, we co-cultured asunaprevir- or vehicle-treated IKAROS-degron cells with mock, blinatumomab, 19.BBz, 19.28z, 22.BBz, and dual 19/22.BBz CAR T cells. At three different effector-to-target ratios, IKAROS^low^ cells were more resilient to CD19- and CD22-targeted therapies (Figures 4D, 4E, 4F, 4G, and 4H, S8), with more pronounced resistance at lower effector-to-target ratios.

**Figure 4:**
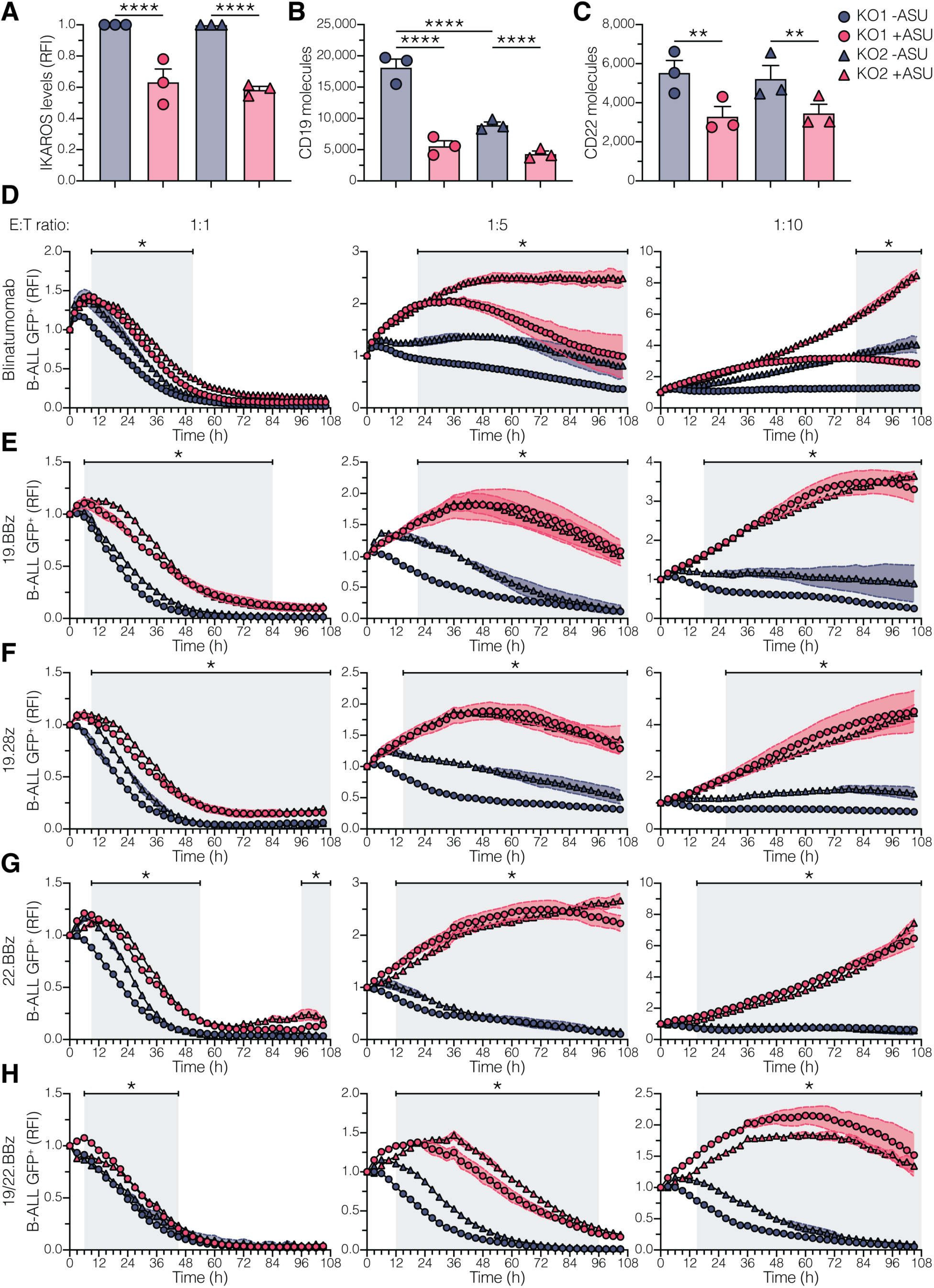
Cells with low IKAROS expression are resistant to CD19- and CD22-targeted therapies. **(A)** Median IKAROS expression in IKAROS-degron models (KO1 and KO2) treated with DMSO (-ASU) or 10 µM asunaprevir (+ASU) for 7 days. Values were measured by flow cytometry and normalized to -ASU condition. RFI = relative fluorescence intensity. **(B – C)** CD19 (B) and CD22 (C) surface quantitation in IKAROS-degron models (KO1 and KO2) treated with DMSO (-ASU) or 10 µM asunaprevir (+ASU) for 7 days. **(D – H)** IKAROS-degron models (KO1 and KO2) were treated with DMSO (-ASU) or 10 µM asunaprevir (+ASU) for 7 days. IKAROS-degron cells were washed out of DMSO or asunaprevir and co-cultured with blinatumomab-treated (D), 19.BBz (E), 19.28z (F), 22.BBz (G), and dual 19/22.BBz (H) CAR T cells at 1:1, 1:5, and 1:10 E:T ratio. B-ALL cell viability was measured at every 2 - 3 h interval via IncuCyte. GFP median values were normalized to 0 h condition. RFI = relative fluorescence intensity. (*) denotes conditions in which both KO1 +ASU *vs.* KO1 -ASU and KO2 +ASU *vs.* KO2 -ASU comparisons were statically significant. Experiments in (A – H) were performed in triplicate with three different T-cell donors. Dots represent the mean value from two (A – C) or three (D – H) technical replicates. Bar plots in (A - C) show mean ± SEM. Curves in (D - H) show mean ± SEM. Statistical tests used were one-way ANOVA followed by Šidák’s multiple comparisons test (A - C); and two-way ANOVA followed by Šidák’s multiple comparisons test (D - H). *P<0.05, **P<0.01, ****P< 0.0001.

To determine if the advantage of IKAROS^low^ cells against these therapies was due to reduced antigen expression and to rule out the effect of other IKAROS-regulated genes, we developed a model whereby CD19 levels were unaffected by IKAROS. We knocked out the endogenous *CD19* gene in our IKAROS-degron models and reintroduced ectopic expression of wild-type CD19. In these models, named CD19KO-FL, asunaprevir treatment reduced IKAROS levels without decreasing CD19 surface expression (Figure S9A), while the changes in other proteins persisted (Figure S9B). As expected, asunaprevir treatment did not provide any advantage against CD19-targeted therapies, even at low E:T ratios (Figures S9C, S9D, and S9E). Thus, decreased CD19 and CD22 surface expression due to low IKAROS levels reduces the efficacy of CD19- and CD22-targeted therapies and B-ALL cell killing, demonstrating that IKAROS^low^ cells have a survival advantage in the face of these therapies.

## Discussion

Antigen loss after targeted immunotherapies, including blinatumomab, CD19- or CD22-directed CAR T cells, remains a significant clinical challenge for patients with r/r B-ALL. CD19^neg^ relapses have also been observed in non-Hodgkin’s lymphoma patients^32,33^. There is no standard of care for patients experiencing a CD19^neg^ relapse, and outcomes are poor^9,34^. Often, CD22 is the next antigen targeted, as it is expressed in the majority of B-ALL^9,10,35,36^. Here, we found that the transcription factor IKAROS modulates CD19 and CD22 surface expression in a dose-dependent and reversible manner. IKAROS is a transcription factor crucial in B cell lineage specification and commitment and is a known B-ALL tumor suppressor^37^. In B-ALL, genetic alterations affecting the IKAROS gene, *IKZF1*, are associated with poor response to front-line therapy and are a factor in relapse risk stratification^29^. However, the role of IKAROS in the failure of CD19- and CD22-targeted therapies has not been described.

Two non-mutually exclusive models explain the source of CD19^neg^ tumor cells. First, the immune-enrichment model posits that rare pre-existing CD19^neg^ tumor cells are selected while antigen-positive cells are eradicated^38,39^. A single-center study of 166 pediatric and young adult r/r B-ALL cases treated with tisagenlecleucel found that the presence of CD19^neg/dim^ tumor cells prior to therapy did not predict nonresponse or recurrence after CART19 therapy^40^. Second, the immune-pressure adaptive model suggests that some CD19^pos^ tumor cells possess intrinsic properties that favor losing CD19 to CD19 targeting^13^. Our data support the latter model. Using single-cell approaches in clinically annotated samples, we found that, prior to CART19 therapy, CD19^pos^ pro-B-like tumor cells with low levels of IKAROS were associated with CD19 loss and CD19^neg^ relapse. Notably, IKAROS levels differed only in pro-B-like cells, not in other subpopulations, highlighting the power of our single-cell approach.

In the present study, we demonstrated that IKAROS^low^ pro-B-like tumor cells are enriched for an HSC/ MPP gene signature, depicting loss of B-cell commitment. CD19 loss has been associated with increased expression of stem/progenitor markers, such as CD34 and CD123, suggesting de-differentiation to a developmentally earlier state^41^. IKAROS KD or regulatable models demonstrated that B-ALL cells acquire the HSC/MPP signature through modulation of IKAROS. Low levels of IKAROS resulted in modulation of B-cell phenotype with gain of stem/progenitor and myeloid proteins while B-lineage proteins decline. These include targets of immune therapies in B-ALL (CD22^42^, CD38^43^, and CD72^44^), suggesting more potential for phenotypic plasticity and lineage infidelity. Wholesale modulation of B-ALL cell state and phenotype aligns with previous observations in cell line models and patients, where CD19 loss was accompanied by CD22 downregulation^18–22^ and loss of both antigens is reversible^17,23^.

Although not addressed in our study, lineage switch relapses occur after CD19-directed therapy^16^. Several studies suggest that the acquisition of myeloid features at relapse originates from CD19^pos^ B-ALL cells through reprogramming or selection of clones with HSPC features^45–48^. Our data suggest that IKAROS^low^ cells have increased plasticity and potential for lineage infidelity and thus may be ripe to support lineage switch. It is likely, however, that only those with the right conditions (genomic background, cytokine signaling) will be able to overcome the barrier to differentiate towards the myeloid compartment, consistent with the model presented by Jacoby et al.^47^

Several mechanisms underlying antigen loss have been described. CD19 loss has been associated with truncating mutations of the *CD19* gene^11^, mutations in genes involved in CD19 membrane trafficking^12,13^, and *CD19* mRNA alternative splicing^14,15^. CD22 downregulation, which is less understood than CD19 loss, has been reported through alterations in transcription and splicing^17^. Further, no evidence suggests that these potential mechanisms are mutually exclusive. We found distinctions in how IKAROS modulates CD19 versus CD22 surface expression. While there were no differences in *CD19* mRNA levels or promotor accessibility in IKAROS KD cells, *CD22* mRNA levels were significantly lower, with changes in the accessibility to two peaks in *CD22* intronic regions. Moreover, the kinetics for CD22 and CD19 surface expression recovery in the regulatable IKAROS model were different. These results suggest that IKAROS modulates CD19 and CD22 surface expression through different mechanisms, which will be the subject of future studies. Despite these distinctions, we show that IKAROS DNA binding domains are required to modulate CD19 and CD22 surface expression. *IKZF1* alterations occur in 25 to 80% of B-ALL cases, with *IKZF1* partial or complete deletions as the main alterations^30^. In our cohort, *IKZF1* deletions were only reported in two patients, and both were refractory to CART19. The presence of *IKZF1* deletions did not correlate with lower *CD19* or *CD22* mRNA or protein levels. Thus, our data suggest that patients with *IKZF1* deletions are not more prone to antigen escape relapse following CD19- or CD22-targeted immunotherapies.

High disease burden prior to treatment, non-response to blinatumomab, and emergence of minimal residual disease are clinical features associated with CD19^neg^ relapse^34,40,49–51^. Prior blinatumomab induces higher expression of alternative CD19 isoforms, decreases CD19 surface expression, and selects tumor cells more fit to survive CD19-directed immunotherapies^13,40,51^. In our CART19 cohort, 5 patients received prior CD19-targeted therapy (blinatumomab or CART19); 4 of them suffered CD19^neg^ relapse, while the other achieved a durable CR. High disease burden may increase the frequency of IKAROS^low^ tumor cells. Furthermore, when challenged with CD19- and CD22-targeted therapies, IKAROS^low^ tumor cells had the greatest survival advantage at lower effector-to-target ratios, conditions that resemble higher tumor burdens, and indicate a higher probability for IKAROS^low^ tumor cells to survive and enable antigen escape relapse.

FDA approval for early administration of blinatumomab in front-line therapy and inotuzumab after the first non-response or relapse increases the population at risk for antigen escape. Hematopoietic stem cell transplantation (HSCT) has been used as a consolidative therapy to avoid antigen escape relapse but still risks long-term complications and side effects^52^. Understanding the mechanisms of antigen loss and thereby identifying patients at risk for antigen escape relapse would change the treatment paradigm for these challenging patients. Our results provide insight into mechanisms of antigen modulation and lineage plasticity mediated by IKAROS dose and demonstrate a potential prognostic role for IKAROS in the early identification of patients at risk of antigen loss and identify IKAROS as a target to revert or subvert these relapses.

## Methods

### Bone marrow samples from patients and healthy donors

Healthy human bone marrow (BM) was purchased from AllCells, Alameda, CA, USA (n = 6; median age was 23 years (range, 21 – 32 years); 5 males and 1 female). De-identified patient-derived xenograft (PDX) samples were obtained from Children’s Hospital of Philadelphia. thirty-nine samples were collected under informed consent from twenty-three patients enrolled in CHP959 (NCT01626495; n = 18), 14BT022 (NCT02228096; n = 4), and 16CT022 (NCT02906371; n = 1) studies; and two adult patients treated compassionately: one patient with r/r B-ALL that underwent CD19^pos^ relapse after CART19 treatment (Patient ID: Pos1) and one patient with chronic lymphocytic leukemia/small lymphocytic lymphoma (CLL/SLL) with Richter’s transformation that underwent CD19^neg^ relapse after CART19 therapy (Patient ID: Neg11). All patients in this cohort were treated with 19.BBz CAR T cells. Twenty-two de-identified primary patient samples were obtained from 11 patients who were consented and treated with 22.BBz CAR T cells on a phase I trial (NCT02315612) at the National Cancer Institute. The Stanford University Institutional Review Boards approved the use of these samples.

### Cell culture and CD19 knock out generation

697, NALM6, NALM16, NALM20, REH, RS4;11, and SUP-B15 cell lines were purchased from ATCC (Manassas, VI, USA). CI, JVM-2, MHH-CALL4, WA-OSEL were purchased from DSMZ (Braunschweig, Germany). OCI-Ly1, OCI-Ly-7, and SUDHL6 were a gift from the Amengual lab^53^. 697, JVM-2, NALM6, NALM16, REH, RS4;11, SUDHL6, and WA-OSEL were cultured in RPMI-1640 medium supplemented with 10% fetal bovine serum (FBS). OCI-Ly1 and OCI-Ly-7 were cultured in IMDM medium supplemented with 10% FBS. CI, MHH-CALL4, NALM20, and SUP-B15 were cultured in IMDM or RPMI-1640 medium supplemented with 20% FBS. For all cell lines, the medium was additionally supplemented with 2 mM L-glutamine (Invitrogen) and 1x penicillin/streptomycin (Invitrogen), and cells were maintained at 37°C and 5% CO_2_.

To generate CD19 KO cell lines, we incubated 3.2 µg synthetic gRNA against the *CD19* gene (Synthego KO kit v2) with 6 µg purified Alt-R S.p. HiFi Cas9 Nuclease V3 (IDT) for 20 min at 37°C. Guide sequences were as follows: 5′-UGCCAGGCCUUCUCAGAGGG-3′, 5′-UUUUAAGAAGGGUUUAAGCG-3′, and 5′-CUUCAACGUCUCUCAACAGA-3′. This ribonucleoprotein complex was transfected into 1.5 x 10^5^ MHH-CALL4, NALM6, REH, or SUP-B15 cells via electroporation using a 4D-Nucleofector system and AMAXA SF reagent with program CV-104 (Lonza). Seven days post-electroporation, cells were processed for CyTOF and CD19^neg^ (CD19 KO) cells were gated from CD19^pos^ (CD19 WT) cells.

### Mass cytometry

Samples were processed as previously described^54^. For viable frozen healthy BM, leukemic primary, and PDX samples, cells were thawed in 90% RPMI medium (ThermoFisher Scientific) with 10% FCS, 20 U/ml sodium heparin (Sigma-Aldrich), 0.025 U/ml Benzonase (Sigma-Aldrich), 1x L-glutamine and 1x penicillin–streptomycin (Invitrogen) and rested at 37 °C for 30 min. For B-ALL cell lines, cells were resuspended in their corresponding culture media. For healthy BM, leukemic primary, and PDX samples, 1 x 10^6^ cells were stained for viability with cisplatin as described^55^. Following viability staining, cells were fixed with 1.6% paraformaldehyde (PFA, Electron Microscopy Sciences) for 10 min at RT. Cells were barcoded using 20-plex palladium barcoding plates prepared in-house as described^56^. We included one healthy BM reference sample within each barcoding plate to control for batch effects. A total of 15 barcode plates were used in this study. Following barcoding, cells were pelleted and washed once with cell-staining medium (CSM; PBS with 0.5% BSA and 0.02% sodium azide) to remove residual PFA. Blocking was performed with Human TruStain FcX (BioLegend), following the manufacturer’s instructions. Antibodies to surface proteins were added, yielding 800-μl final reaction volumes, and samples were incubated at RT and 300 rpm for 30 min. Cells were washed with CSM before permeabilization with methanol for 10 min at 4°C. Cells were washed with CSM and stained with intracellular protein and phospho-specific antibodies in 800 μl for 30 min at RT and 300 rpm. Cells were washed once in CSM, then stained overnight with 1:5,000 191Ir/193Ir or 103Rh DNA intercalator (Standard Biotools) in PBS with 1.6% PFA at 4°C. Cells were washed once with CSM, washed twice with double distilled water, filtered to remove aggregates, and resuspended in 139La/ 142Pr/ 159Tb/ 169Tm/ 175Lu normalization beads^57^ immediately before analysis using a Helios mass cytometer (Standard Biotools). Throughout the analysis, cells were maintained at 4°C and introduced at a constant rate of 150 - 200 cells/s.

### Processing of mass cytometry data

Data were normalized together using bead normalization^57^ and files were debarcoded as described^56^. Single-cell protein expression data were extracted and analyzed using packages from the Comprehensive R Archive Network (CRAN) project (https://cran.r-project.org/) and Bioconductor (http://www.bioconductor.org). Raw data were transformed using the inverse hyperbolic sine (arcsinh) function with a cofactor of 5. Expression of proteins in each population of interest was determined by calculating the median level of expression after arcsinh transformation. For UMAP visualization, each population or sample was randomly subsampled to 1,000 cells. Dimensionality reduction was performed using umap package (version 0.2.9.0) based on arcsinh values of selected markers.

### Manual gating

Single cells were gated using Community Cytobank software (https://community.cytobank.org/) based on event length and 191Ir/193Ir or 103Rh DNA content (to avoid debris and doublets) as described^58^. For B-ALL cell lines, live B-ALL cells were gated based on cleaved poly(ADP-ribose) polymerase (cPARP) and ^195^Pt or cleaved Caspase3 (cCaspase3) content^55^. Downstream analyses were performed in this live B-ALL cell fraction. For healthy BM, leukemic primary, and PDX samples, following single-cell gating, live non-apoptotic cells were gated based on cleaved poly(ADP-ribose) polymerase (cPARP) and ^195^Pt content^55^. In PDX samples, murine cells were excluded by gating on mouse CD45 (mCD45) protein. Platelets and erythrocytes were excluded by gating on CD61 and CD235a/b, respectively. The remaining fraction was gated to exclude T cells and myeloid cells based on CD3e, CD11b, CD16, and CD33 expression. After further exclusion of CD38^high^ plasma cells, the remaining fraction was defined as lineage-negative blasts (Lin−/ B+; see Figure S2A for gating). Further analyses were applied to Lin−/ B+ fraction. For the CART22 cohort, since some samples had less than 1% blast cells, we applied opt-SNE^59^ to the Lin-/B+ fraction of each paired sample together with their corresponding healthy BM reference to gate blast cells, as was previously shown^60^. The following markers were used for opt-SNE: CD10, CD11b, CD16, CD179b, CD24, CD33, CD34, CD38, CD45, CD81, HLADR, intracellular immunoglobulin heavy chain (IgMi), surface immunoglobulin heavy chain (IgMs), Ki67, phosphorylated CREB, phosphorylated S6, and terminal deoxynucleotidyl transferase (TdT).

### B-cell developmental classification

We used the single-cell developmental classifier previously reported^25^. Briefly, Lin−/ B+ fraction from healthy human BM was gated into 15 developmental populations of normal B lymphopoiesis, mixed progenitors, and mature non-B cell fractions, as shown in Figure S1. The distribution of each population was based on the expression of 10 B cell developmental proteins that were used for manual gating: CD19, CD20, CD24, CD34, CD38, CD45, CD127, CD179b, IgMi, and TdT. Lin−/ B+ cells from each leukemia sample or live B-ALL cells from cell lines samples were assigned to the most similar healthy fraction based on the shortest Mahalanobis distance among distances to all healthy developmental populations in these ten dimensions. A cell was designated ’unclassified’ if none of the distances were below the classification threshold (Mahalanobis distance = 10, based on the number of dimensions).

### Single cell RNA and antibody tag sequencing

Viably frozen healthy BM and PDX samples (Patient ID: CR2, CR4, CR6, Neg2, Neg5, Neg11) were thawed in 90% RPMI medium (ThermoFisher Scientific) with 10% FCS, 20 U/ml sodium heparin (Sigma-Aldrich), 0.025 U/ml Benzonase (Sigma-Aldrich), 1x L-glutamine and 1x penicillin–streptomycin (Invitrogen) and rested at 37 °C for 30 min. Then cells were filtered through cell strainer size 35 µm and centrifuged at 350g for 5 min. Cells were resuspended in 1 ml of Stain Buffer (BD Biosciences) and blocking was performed with Human Fc Block (BD Biosciences) following the manufacturer’s instructions. To enrich for Lin−/ B+ fraction, samples were incubated with biotin-conjugated antibodies for 30 min. Cells were washed with Stain Buffer and then incubated with BD Streptavidin Particles Plus (BD Biosciences) at the manufacturer’s recommended concentration for 30 min at RT. Particle-labeled cells were placed in a magnetic holder for 6 min. The supernatant was transferred to a new tube and placed back in the magnetic holder for an additional round of depletion and supernatant recovery. Cells from the supernatant were then pelleted by centrifugation at 350g for 5 min.

Lin-/ B+ fraction was resuspended in 180 µl of Stain Buffer supplemented with a mix of BD AbSeq oligo conjugated antibodies against CD19, CD20, CD24, CD34, CD38, CD45, CD127, and IgM (BD Biosciences). Then, each sample was labelled with the Human Single Cell Sample Multiplexing kit (BD Biosciences) and incubated at RT for 30 min. Cells were washed twice with Stain Buffer and resuspended in Sample Buffer (BD Biosciences). Cell number was counted with Countess II Cell Counter (Thermo Fisher Scientific). A total of 50,000 cells (12,500 cells per sample, up to 4 samples) were pooled together, and single cells were isolated in a BD Rhapsody cartridge using BD Rhapsody Express Single-Cell Analysis System (BD Biosciences). We included one healthy BM reference sample within each cartridge to control for batch effects. A total of 3 cartridges were used in this study. For each cartridge, we followed the manufacturer’s instructions to prepare whole transcriptomic, antibody tag, and sample tag libraries using the Whole Transcriptome Analysis (WTA) Amplification Kit (BD Biosciences). Libraries from the same cartridge were indexed with identical Illumina sequencing adapters. Final libraries were pooled together sequencing on a NovaSeq 6000 sequencer (Illumina) at MedGenome (Foster City, CA, USA) with paired-end 100 base pair (bp) reads.

### Processing of CITE-seq data

Fastq files were processed through the Rhapsody analysis pipeline (BD Biosciences) on the Seven Bridges platform (https://www.sevenbridges.com) following the manufacturer’s recommendations. Reads were mapped to the hg38 reference genome using bowtie2. Final expression matrices contain recursive substation error correction (RSEC) adjusted molecule counts per cell in a CSV format. Molecule count tables were read into the R package Seurat (version 3.2). Out of 35,716 total cells, 2,106 (5.9 %) were multiplets, and 637 (1.8 %) were undetermined events. The mean number of cells sequenced per sample was 2,748. The mean number of genes detected per cell was 947, with a mean of 2,670 reads/cell for the mRNA library and 1,493 reads/cell for the antibody tag library. Cells with less than 500 expressed genes, or 200 gene reads assigned, and more than 50 % of genes being mitochondrial genes were excluded for downstream analysis, removing 12,597 cells. We normalized transcriptomic data through variance stabilizing transformation with the removal of mitochondrial gene percentage as a potential confounding source of variation. Antibody tag data was normalized with a centered log-ratio transformation.

We performed the Wilcoxon rank sum test for differential expression analysis followed by Bonferroni correction. Genes and antibody tags expressed in at least 10 % of cells from one of the conditions under comparison, had an absolute fold change higher or equal to 25%, and had adjusted p-value lower than 0.05 were called significant. Heatmaps were plotted using the ComplexHeatmap package (version 2.6.2), and the mean expression of each gene or antibody tag per population. To estimate the single-cell enrichment score for the HSC multipotent progenitor program, we used the function AddModuleScore from the Seurat package.

### Projection of tumor cells onto healthy BM UMAP space

To define healthy BM populations and their UMAP space, we performed principal component (PC) analysis on 3,000 most variable genes (MVG) across all healthy BM cells and antibody tag data. Based on the top 30 PCs, we performed UMAP embedding of healthy cells, as well as unsupervised clustering with the functions FindNeighbors and FindClusters from the Seurat package. We used the function FindMarkers to identify genes significantly up-regulated in each cluster against all the other clusters. Differentially expressed genes were analyzed with Enrichr^61^ (https://maayanlab.cloud/Enrichr/) to define biological pathways associated with each cluster and assign their corresponding cell population.

To project tumor cells onto the healthy BM space, we predicted their top 30 PCs based on the PC analysis performed for the healthy BMs and the expression of 3,000 MVG defined previously and antibody tag data. These predicted top 30 PCs were used to project and embed tumor cells onto healthy BM UMAP space. To associate each tumor cell with their closest healthy population, we used a k-nearest neighbors model (k-NN). Briefly, we performed 10-fold cross-validation to train a k-NN model to predict the cell population of healthy BM cells based on the top 30 PCs. For this model, the ten nearest neighbors were considered, and the decision was taken based on majority voting. Finally, we defined the closest population to each tumor cell by applying this k-NN model to their predicted top 30 PCs.

### Retrovirus and Lentivirus production

For retrovirus production, 293GP cells were used (graciously provided by Dr Garry Nolan). Briefly, 70% confluent 293GP on 10-cm poly-d-lysine coated plates were cotransfected with 9 μg of IKAROS-degron-P2A-mNeonGreen, IK6-degron-P2A-mNeonGreen, CD19WT-IRES-eGFP, 19.BBz, 19.28z, 22.BBz, or 19/22.BBz CAR encoding vectors and 4.5 μg RD114 envelope plasmid (graciously provided by Dr. Crystal Mackall) with lipofectamine 3000 (Invitrogen). Viral supernatants were collected 48 and 72 hours post-transfection, centrifuged to separate cell debris from the viral supernatant, and frozen at −80°C for future use.

For third-generation, self-inactivating lentivirus production, 4 – 5 × 10^6^ 293T cells (graciously provided by Dr. Crystal Mackall) were plated on a 10-cm dish for 24 hours before transfection. On the day of transfection, a mixture of 1.5 µg pMD2.G envelope plasmid (Addgene #12259), 2.25 µg psPAX2 packing plasmid (Addgene #12260), and 3.75 µg vector plasmid were cotransfected with TurboFect (Thermo Fisher). For the current study, we used the following vector plasmids: pLKO-RFP-shCntrl (Addgene #69040), pLKO-RFP-IKZF1-sh2 (Addgene #69041), pLKO-RFP-IKZF1-sh3 (Addgene #69042), SMARTvector Inducible Non-targeting PGK-TurboRFP (VSC11656, Horizon Discovery Biosciences), and SMARTvector Inducible Human IKZF1 PGK-TurboRFP shRNA 1 (V3SH11252-224727699, Horizon Discovery Biosciences). 48 and 72 hours post-transfection, the supernatant was collected and centrifuged to separate cell debris from the viral supernatant. PEG-it solution (System Biosciences Innovation) was added to the viral supernatant, and lentivirus particles were concentrated following the manufacturer’s protocol. Concentrated lentivirus aliquots (20 µl) were stored at −80°C.

### Lentiviral and retroviral transduction

For lentivirus transduction, 1 × 10^6^ cells were incubated with 20 µl of concentrated lentivirus particles and TransDux Max reagents (System Biosciences Innovation) following the manufacturer’s instructions. 48 hours post-transduction, cells were washed and incubated in their corresponding complete media. When appropriate, 1 µg/ml puromycin (Invivogen) was added to the complete media for antibiotic selection and 250 ng/ml doxycycline (Sigma) for expression of doxycycline-inducible systems. Transduction efficiency was followed by measuring RFP expression in a CytoFLEX instrument (Beckman Coulter).

For retroviral transduction, non-tissue culture treated 12-well plates were coated overnight at 4°C with 25 μg/ml retronectin in PBS (Takara). Plates were washed with PBS and blocked with PBS supplemented with 2% BSA for 15 min. Thawed retroviral supernatant was added at ∼1 ml per well and centrifuged at 3,200 rpm for 1.5 hours before adding 0.5 × 10^6^ cells. After 48 hours, transduction efficiency was checked by GFP expression in CytoFLEX instrument (Beckman Coulter). When required, transduced cells were enriched by sorting GFP-positive cells in a BD FACSAria II SORP Cell Sorter (BD Biosciences).

### Lenalidomide treatment

For targeted degradation of IKAROS protein, 5 x 10^5^ tumor cells were seeded in duplicate in a 24-well plate with 2 ml complete media supplemented with 0 µM, 0.1 µM, 1 µM, or 10 µM lenalidomide (Selleck Chemicals). DMSO was used as a vehicle to adjust the drug volume added. After 3, 4, or 7 days, CD19 and IKAROS levels were measured by flow cytometry.

### Flow cytometry analysis

To assess CD19 and CD22 surface expression, 0.5 – 1 x 10^5^ cells were washed with CSM and incubated with 100 µl antibody mix (1 µl APC anti-human CD19 (clone: HIB19, BioLegend); or 1 µl APC anti-human CD22 (clone: HIB22, BioLegend) with or without 1 µl Pacific Blue anti-human CD19 (clone: HIB19, BioLegend) in 100 µl CSM) at RT for 10 min. Cells were washed twice and resuspended in 200 µl CSM for flow analysis in CytoFLEX cytometer (Beckman Coulter).

To assess IKAROS intracellular levels, 0.5 – 1 x 10^5^ cells were washed with CSM and fixed with 1.6% PFA (Electron Microscopy Sciences) in CSM for 10 min at room temperature (RT). Cells were washed twice with CSM and incubated with 1 µl Pacific Blue anti-human CD19 antibody (clone: HIB19, BioLegend) in 100 µl CSM at RT for 10 min. Cells were washed twice and permeabilized with 100 µl Methanol at 4°C for 10 min. After three washes, cells were incubated with 1 µl Alexa Flour 647 anti-human IKAROS antibody (clone: 16B5C71, BioLegend) in 100 µl CSM at RT and 300 rpm for 30 min. Cells were washed twice and resuspended in 200 µl CSM for flow analysis in CytoFLEX cytometer (Beckman Coulter).

Quantification of CD19 and CD22 surface molecules on cancer lines was performed using the BD Quantibrite™ APC Fluorescence Quantitation Kit (BD Biosciences).

Flow cytometry data was analyzed using Community Cytobank software (https://community.cytobank.org/). Briefly, forward versus side scatter was used to exclude debris, while forward scatter area versus width was used for doublet exclusion. When pertinent, tumor cells were gated based on GFP (IKAROS-degron, IKAROS-degron CD19KO-FL, IK6-degron) or RFP (IKAROS WT and KD cells) expression. Finally, CD19, CD22, and IKAROS median fluorescent intensities (MFI) were calculated in the gated population.

### Generation IKAROS KD cells for ATAC-seq and RNA-seq

5 x 10^6^ B-ALL cells (NALM6, REH or SUP-B15) were transduced with lentivirus expressing *IKZF1* specific (Addgene #69041) or scramble (Addgene #69040) shRNA in duplicate. 72 hours post-transduction, 1 x 10^5^ cells were used to check transduction efficiency, CD19 and IKAROS levels via flow cytometry analysis as described. In all cases, transduction efficiencies were over 90%.

For ATAC-seq, 50,000 cells were processed as previously reported^62^. For RNA-seq, total RNA was extracted from 3 – 5 x 10^6^ cells using Rneasy kit with Dnase I treatment (QIAGEN), following manufacture instructions. RNA libraries were performed with TruSeq stranded mRNA kit (Illumina). ATAC and RNA libraries were sequenced on the Illumina NovaSeq 6000 at MedGenome facility with paired-end 50 bp or 100 bp reads, respectively.

### RNA-seq data processing

For isogenic IKAROS WT or KD B-ALL cells RNA-seq data, we sequenced an average of 42 x 10^6^ reads per sample (range: 29 x 10^6^ – 60 x 10^6^ reads). Paired-end reads were aligned and quantified using Salmon (version 1.2.0) index against the hg38 reference genome. Gencode transcript annotations (version 37) were used for the genomic location of transcriptomic units. Reads aligning to annotated regions were summarized as counts using the R package tximport (version 1.18.0). Differential expression analyses between IKAROS WT and KD samples were performed using DESeq2 (version 1.30.1)^63^. A false discovery rate (FDR) cutoff of 0.05 was used for gene selection. Read counts were normalized using variance-stabilizing transformation (vst) built into the DESeq2 package. Gene set enrichment analysis (GSEA) was performed using the GSEA software (Broad Institute).

Raw fastq files from diagnosis or relapse B-ALL patients (n = 282) from the Therapeutically Applicable Research to Generate Effective Treatments (TARGET) initiative were processed as described above.

### ATAC-seq data processing

ATAC-seq data was processed as previously described^64^. Briefly, adapters were trimmed using cutadapt, and reads were mapped using bowtie2 with a max fragment length of 2000 bp to hg38. We then filtered for non-mitochondrial reads, mapq > 20, and properly paired reads, and removed duplicates using Picard tools. De-duplicated libraries were down-sampled to 15 x 10^6^ read pairs. Aligned, de-duplicated bam files were loaded into R using DsATAC.bam function in the ChrAccR R package. Peaks were called using macs2 with the following parameters on Tn5 insertion sites: - -shift -75 –extsize 150 –nomodel –call-summits –nolambda -p 0.01 -B –SPMR. The consensus peak set across technical and biological replicates was calculated using the getConsensusPeakSet function in the ChrAccR R package. The count matrix was calculated as insertion counts across samples at consensus peak set regions using the ChrAccR regionAggregation function. DESeq2^63^ was used to calculate differentially accessible peaks. Differentially accessible peaks were used to calculate motif enrichment (getMotifEnrichment function in the ChrAccR R package) using the human_pwms_v2 motif database (from the chromVARmotifs package). Adjusted p value (FDR value) was converted to -log(FDR value), and top enriched motifs were plotted by -log(FDR value). Genomic localization of consensus peak set was identified using the annotatePeak function in the ChIPseekerR package and was visualized using the IGV browser^65^. Genes harboring differentially accessible peaks on their promotor region (-3000 to + 3000 bp from transcriptional starting site (TSS)) were used for pathways enrichment analysis in Enrichr (https://maayanlab.cloud/Enrichr/). We used Venny 2.1 (https://bioinfogp.cnb.csic.es/tools/venny/) to define up-regulated genes that had higher accessibility to their promoter in IKAROS KD cells. Protein-protein interaction network between these genes was created with STRING^66^ (https://string-db.org/), and Enrichr (https://maayanlab.cloud/Enrichr/) was used to define their lineage associations.

### Viral vector construction

All DNA constructs were visualized using SnapGene software (v.7.2.1; Dotmatics), and cloning was performed using In-fusion seamless cloning (Takara). IKAROS-degron, IK6-degron, and CD19WT sequences were ordered from IDT. IKAROS-degron and IK6-degron were cloned into a XhoI/EcoRI-digested mNeonGreen – firefly luciferase vector (graciously provided by Dr. Crystal Mackall)^67^. CD19WT was inserted into SrfI/XhoI digested pMSCV-FLAG-hIKAROS-IRES-GFP vector (Addgene 74046).

19.BBz and 19.28z CAR constructs were previously cloned into the MSGV1 retroviral vector^67^. In brief, both constructs consisted of the FMC63 scFv, with 19.BBz comprising the sequence FMC63 scFv - CD8 hinge - CD8 transmembrane - 4-1BB costimulatory domain - CD3z and 19.28z comprising the sequence FMC63 scFv - CD28 hinge - CD28 transmembrane - CD28 costimulatory domain - CD3z. The 22.BBz and 19/22.BBz constructs were generated by replacing the FMC63 scFv of the 19.BBz sequence. For 22.BBz, an scFv derived from the m971 antibody was used. For 19/22.BBz, the following sequence was used: GMCSF leader sequence - FMC63 VH - G4S linker - m971 VL - Whitlow linker - m971 VH - G4S linker - FMC63 VL.

### IKAROS-degron and IK6-degron tumor models

5 x 10^5^ REH cells were transduced with IKAROS-degron-P2A-mNeonGreen or IK6-degron-P2A-mNeonGreen expressing retrovirus. After three days, transduced cells were sorted based on GFP expression using a BD FACSAria II SORP Cell Sorter (BD Biosciences). Cells were expanded for seven days and knocked out endogenous *IKZF1*gene using the following synthetic gRNA guides (Synthego KO kit v2): 5′-UGUCGUAGGGCGUGUCGGAC-3′, 5′-CAACAACGCCAUCAACUACC-3′, and 5′-ACCACCUCGGAACCGCCCGG-3′, and the same protocol as described above. Cells were expanded for seven more days before single-cell cloning by limiting dilution into 96-well plates. Wells containing cells were grown to dense cultures before analysis of endogenous *IKZF1* locus knockout efficiency using ICE CRISPR Analysis Tool (Synthego). Clones with 100% knockout efficiency were further characterized by IKAROS, CD19, and CD22 response to asunaprevir treatment.

To generate IKAROS-degron CD19KO-FL tumor models, endogenous *CD19* locus was knocked out in the IKAROS-degron models, as described above. After seven days, CD19^neg^ IKAROS-degron cells were sorted using a BD FACSAria II SORP Cell Sorter (BD Biosciences). Cells were expanded for seven more days and then transduced with CD19WT-IRES-eGFP. After three days, transduced cells were sorted based on CD19 expression using a BD FACSAria II SORP Cell Sorter (BD Biosciences).

### Asunaprevir treatment

5 x 10^5^ IKAROS-degron, IK6-degron, and IKAROS-degron CD19KO-FL tumor cells were seeded in duplicate in a 24-well plate with 2 ml complete media supplemented with 0 µM, 0.1 µM, 1 µM, or 10 µM asunaprevir (Selleck Chemicals). DMSO was used as a vehicle to adjust the drug volume added. After 3, 7, or 22 days, CD19, CD22, and IKAROS levels were measured by flow cytometry. When indicated, 1 x 10^6^ cells were processed for CyTOF, as described above.

For expression recovery experiments, 5 x 10^5^ IKAROS-degron tumor cells were treated with 10 µM asunaprevir or DMSO for 7 days. After that, asunaprevir-treated cells were washed, and one-half were cultured in completed media supplemented with DMSO (ASU/DMSO condition), while the other half remained in media supplemented with 10 µM asunaprevir (ASU/ASU condition). DMSO-treated cells were maintained in media supplemented with DMSO (DMSO condition). After 9, 14, 21, and 28 days, CD19, CD22, and IKAROS levels were measured by flow cytometry.

### CAR T cell production

Leukopaks from healthy donors were purchased from StemCell Technologies. Primary human T cells were purified by negative selection using the RosetteSep Human T cell Enrichment kit (StemCell Technologies) and SepMate-50 tubes. T cells were cryopreserved at 2 × 10^7^ cells per ml in CryoStor CS10 cryopreservation medium (StemCell Technologies) until use. Isolated T-cells were activated and transduced with 19.BBz, 19.28z, 22.BBz, or 19/22.BBz CARs expressing retrovirus as described previously^67^. Unmanipulated (Mock) cells were used as control. CAR T-cell killing *in vitro* assays were performed on day 10 after activation.

### IncuCyte tumor killing assays

IKAROS-degron and IKAROS-degron CD19KO-FL models were treated for 7 days with 10 µM asunaprevir or DMSO. On Day 0, tumor cells were washed and set in co-cultured with mock, blinatumomab-treated (50 ng/ml, Invivogen), 19.BBz, 19.28z, 22.BBz, or 19/22.BBz CAR T cells. 5 × 10^4^ GFP-labelled tumor cells was cocultured with 5, 1, or 0.5 × 10^4^ CAR T cells (E:T 1:1, 1:5, 1:10) in 200 μl RPMI supplemented with 10% FBS, 10 mM HEPES, 2mM l-glutamine, 100 U ml−1 penicillin and 100 μg ml−1 streptomycin. Triplicate wells were plated in 96-well flat-bottom plates for each condition. Tumor fluorescence was monitored every 2–3 h with a ×10 objective using the IncuCyte S3 Live-Cell Analysis System (Sartorius), housed in a cell culture incubator at 37 °C and 5% CO2, set to take 4 images per well at each timepoint. The total integrated GFP intensity was quantified using the IncuCyte basic analyzer software feature (IncuCyte S3 v.2019B Rev2; Sartorius). Data were normalized to the first timepoint and plotted as the fold change in tumor fluorescence over time.

### Statistical analysis

Data were analyzed and visualized using R statistical software (http://www.r-project.org) or GraphPad Prism software. The P values were calculated with the statistical test described in the relevant figure legend. P < 0.05 was considered statistically significant, and P values are denoted with asterisks as follows (P > 0.05, not significant, n.s.; *, P < 0.05; **, P < 0.01; ***, P < 0.001; and ****, P < 0.0001).

## Acknowledgments

We thank current and past members of the Davis laboratory, Ruth Wang’ondu for stimulating discussions, and Joseph Musmacker, Daniel Silberman, and Adam Zoubeidi (BD Biosciences) for technical support in generating and processing CITE-seq data. This work was supported in part by the Intramural Research Program, Center of Cancer Research, National Cancer Institute and NIH Clinical Center, National Institutes of Health (N.N.S., ZIA BC 011823), the V Foundation for Cancer Research, and the National Institutes of Health R01-CA251858. P.D. is supported in part by the American Society of Hematology (ASH) Scholar Award. J.S. is supported by Associazione Italiana per la Ricerca sul Cancro (AIRC, grant no. 27325). N.N.S. receives research funding from Lentigen, VOR Bio, and CARGO Therapeutics. N.N.S. has attended advisory board meetings (no honoraria) for VOR, ImmunoACT, and Sobi. K.L.D is supported in part by MCHRI as the Anne T. and Robert M. Bass Endowed Faculty Scholar in Pediatric Cancer and Blood Diseases. K.L.D has received honoraria and participated in advisory boards for Novartis, and received research funding from Jazz Pharmaceuticals. FACSAria II SORP Cell Sorter (BD Biosciences) was purchased using a NIH S10 Shared Instrumentation Grant (1S10RR02933801). The content of this publication does not necessarily reflect the views of policies of the Department of Health and Human Services, nor does mention of trade names, commercial products, or organizations imply endorsement by the U.S. Government.

## Author contributions

P.D. led this study, designed and performed experiments, analyzed and interpreted data, generated the figures, and wrote the manuscript. J.S. and A.J.M. contributed to mass cytometry experiments and to validate the antibody panel used in these experiments. M.M. and Y.L. contributed with cell culture work, retrovirus, and lentivirus generation and transduction, and flow cytometry data acquisition. K.Z.P, M.C.R., and S.A.Y-H. generated 19.BBz, 19.28z, 22.BBz, and 19/22.BBz CAR T cells and performed IncuCyte experiments. W.D.R. performed ATAC-seq libraries. B.S. and S.C.B. provided guidance for ATAC-seq experiments. R.B. contributed with ATAC-seq data analysis. B.J.S. and A.A.A. provided clinical and RNA-seq data. C.G.M., A.B.L., R.M.M, S.A.G., B.Y., H.W.W., N.N.S, and D.M.B. contributed patient samples and provided clinical data on the patients. J.S., M.C.R., R.G.M., C.L.M., D.M.B., and E.S. provided guidance and scientific input. K.L.D. conceived of and designed this study, provided guidance and scientific input, interpreted data, and wrote the manuscript. All authors discussed the results and commented on the manuscript.

## Supplemental information

**Supplementary Figure S1:**
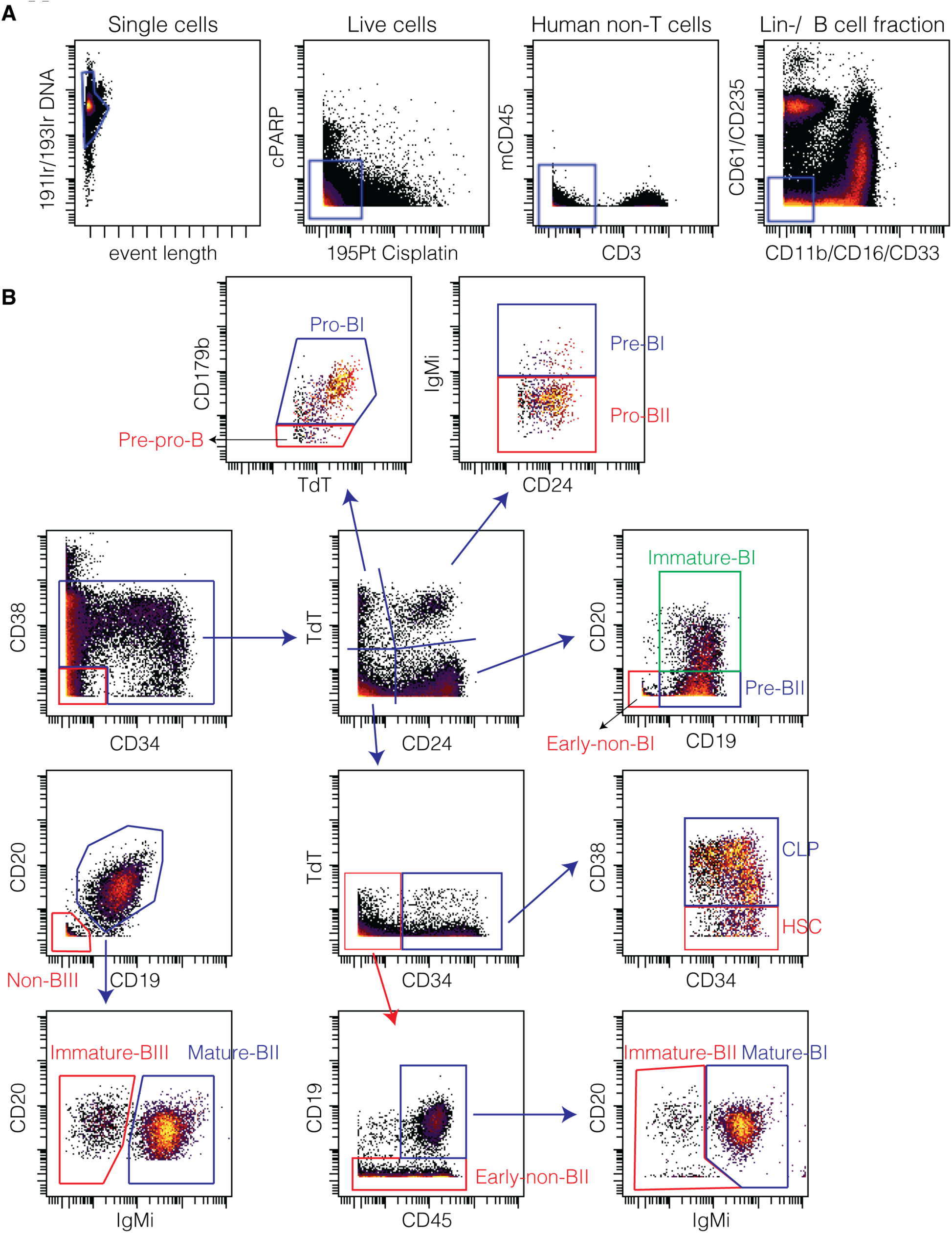
Manual gating strategy. **(A)** Gating strategy for lineage-negative/ B cell (Lin-/ B+) fraction enrichment from healthy BM or leukemia cells. This is the starting population for all subsequent analyses of normal and leukemic samples. (**B**) Gating strategy to identify 12 subpopulations of B lymphopoiesis and 3 non-B populations among Lin-/ B+ fraction from healthy BM.

**Supplementary Figure S2:**
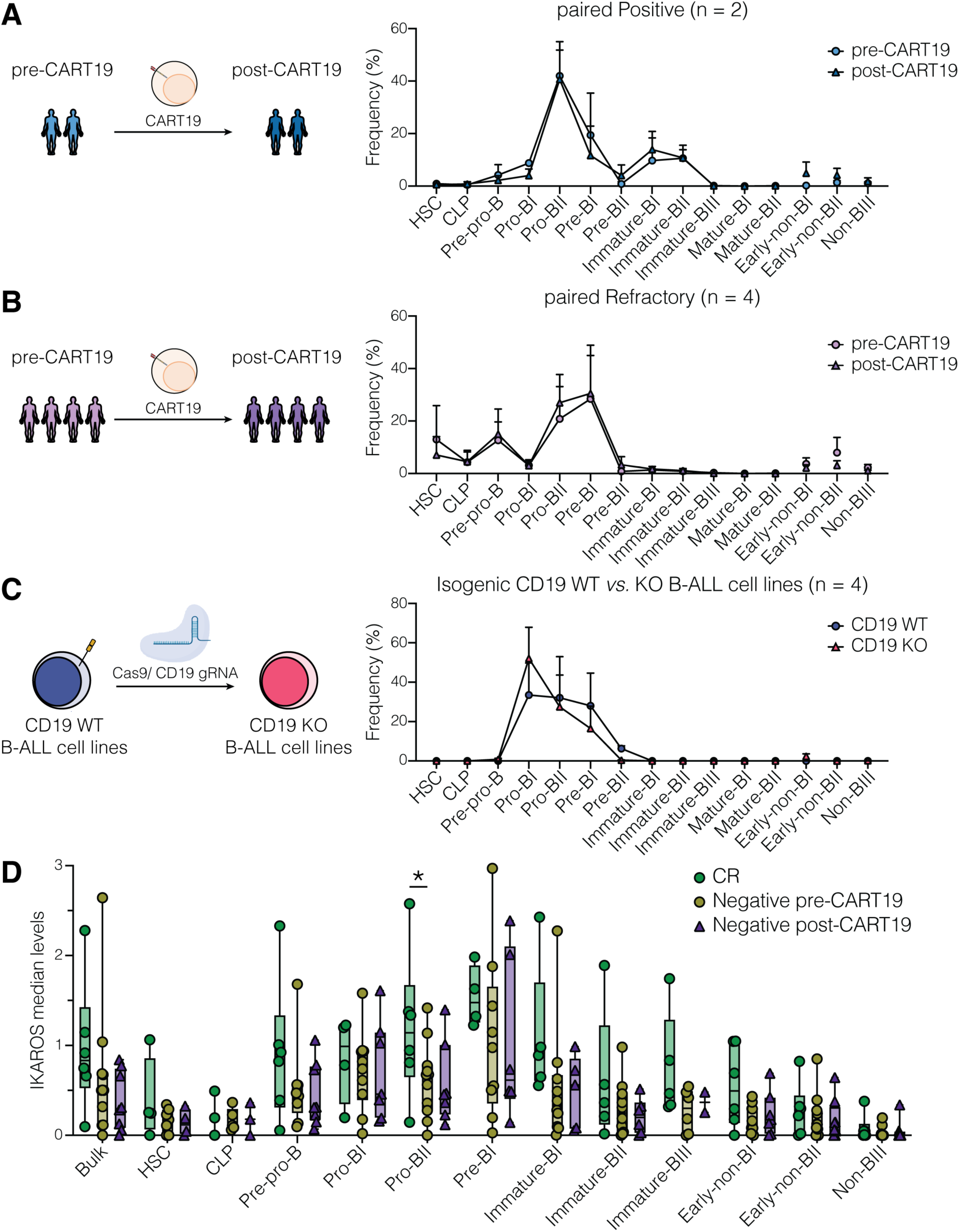
Enrichment in early-non-BI population is restricted to CD19^neg^ relapse patients. **(A - C)** Developmental classification of pre-CART19 and paired post-CD19^pos^ relapse (A; n = 2), refractory (B; n = 4) samples, and isogenic CD19 WT or KO B-ALL cell lines (C; n = 4). **(D)** IKAROS median levels in bulk and different B-cell developmental populations from pre-CART19 patients that achieved durable CR (n = 6), suffered CD19^neg^ relapse (n = 11), and post-CD19 loss (n = 8), respectively. Curves in (A - C) show mean ± SEM. Boxplots in (D) show median with range from minimum to maximum values. Statistical tests used were paired two-way ANOVA followed by Šidák’s multiple comparisons tests (A - C); and two-way ANOVA followed by Tukey’s multiple comparisons tests (D). *P<0.05.

**Supplementary Figure S3:**
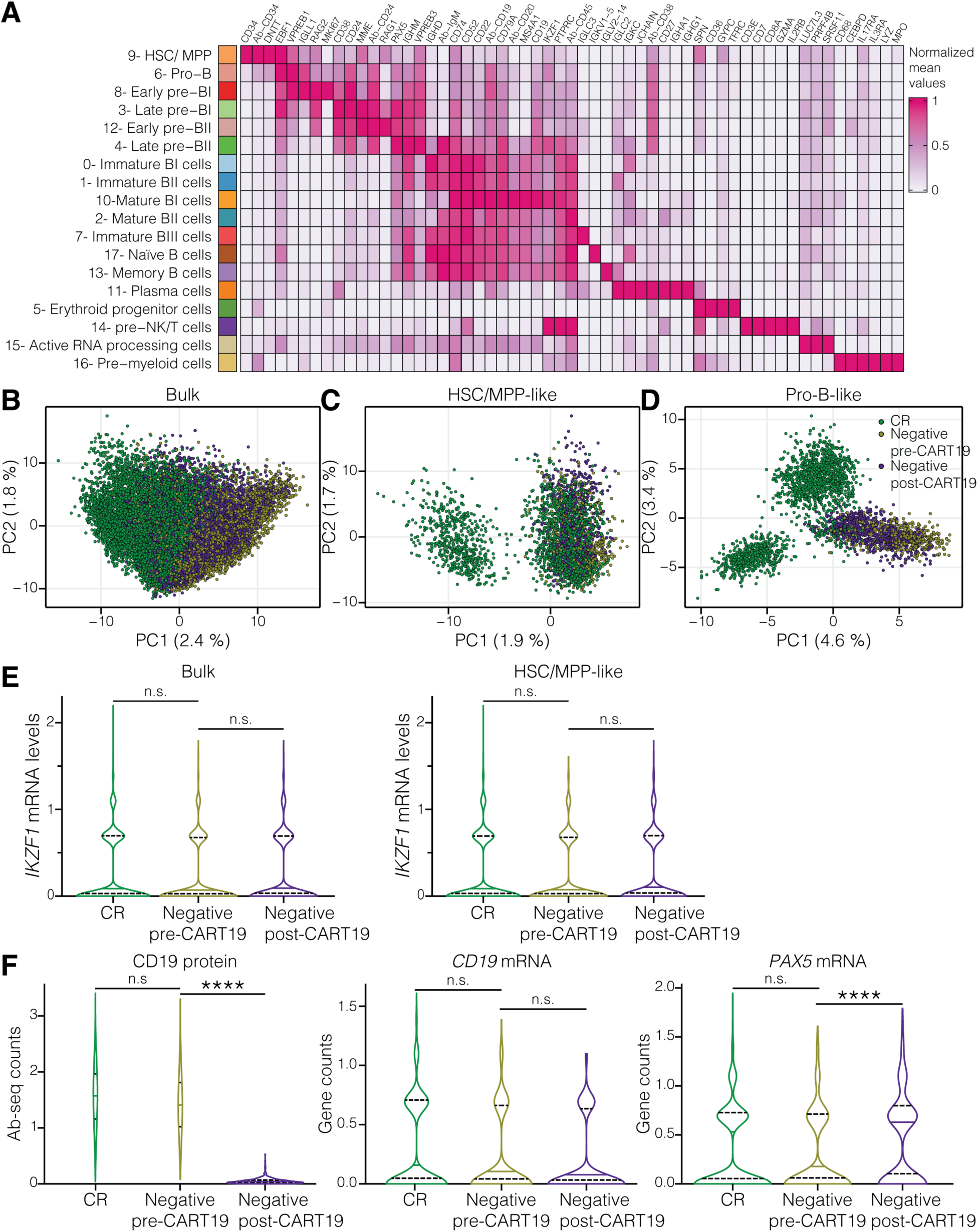
Pre-treatment low *IKZF1* expression and unique gene expression signature in pro-B-like B-ALL cells from CD19^neg^ relapsed patients. (**A**) Mean expression of selected significant markers (FDR < 0.05) for each healthy cluster. Differential expression analysis (Wilcoxon Rank Sum test) was performed between cells from one cluster against cells from all the other clusters. Mean values of each marker were scaled from 0 to 1, to facilitate the visualization of changes in marker expression across different populations. (**B** - **D).** Principal component analysis based on genes differentially expressed between bulk (B), HSC/ MPPC-like (C), and pro-B-like (D) B-ALL cells from pre-CART19 patients that will achieve durable CR or suffer CD19^neg^ relapse. **(E)** *IKZF1* gene expression in bulk (left) and HSC/MPP-like (right) B-ALL cells. (**F**) CD19 protein (left), *CD19* (middle), and *PAX5* (right) gene expression in pro-B-like B-ALL cells. Violin plots in (E - F) show median (solid line) and 25^th^ and 75^th^ quantile (dash lines). Statistical test used was Wilcoxon rank sum test followed by Bonferroni’s multiple comparisons test (E). ****P< 0.0001; not significant (n.s.), P>0.05.

**Supplementary Figure S4:**
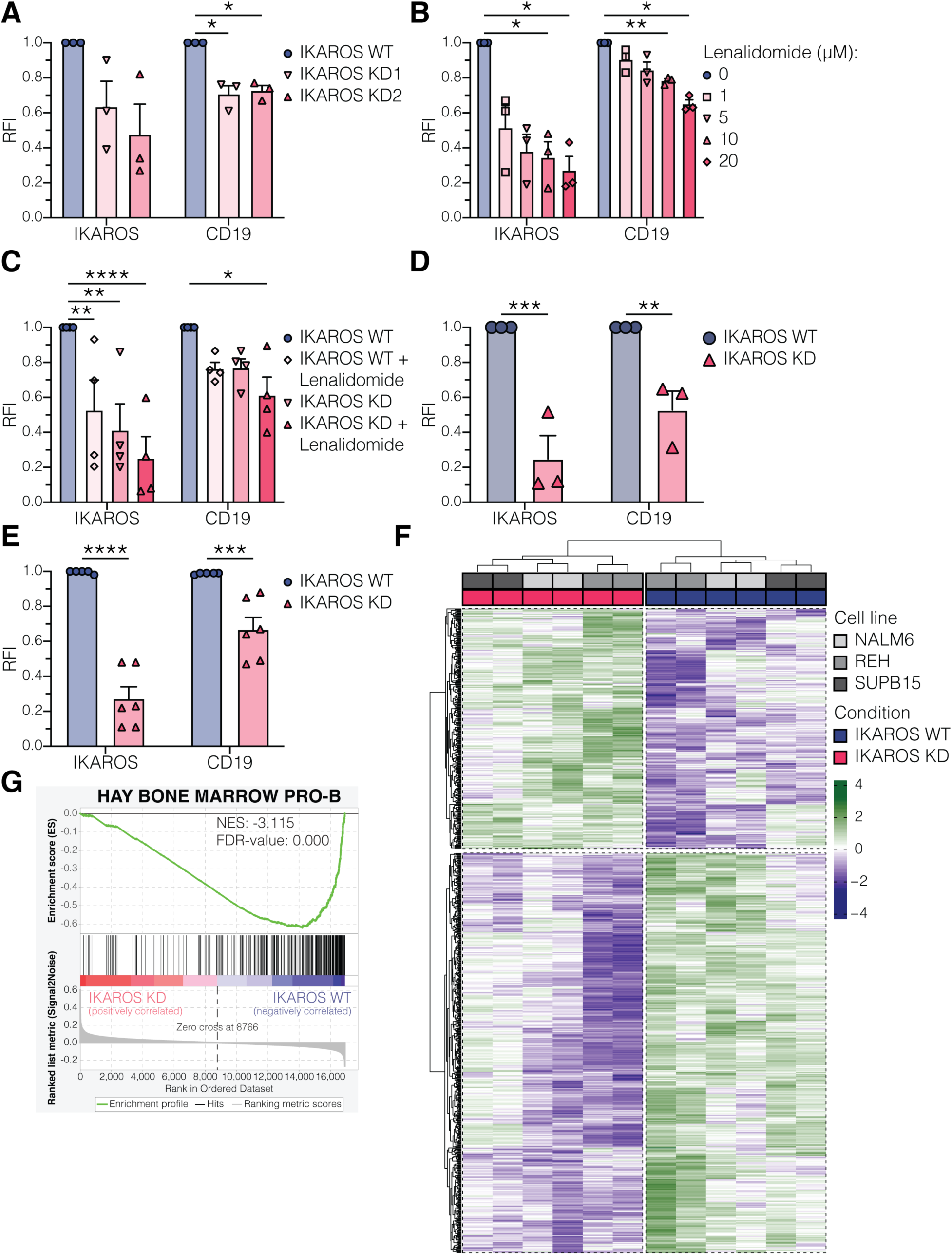
IKAROS modulates CD19 surface expression in other B-cell malignancies. (**A - C**) IKAROS and CD19 median expression values in LBCL cell lines (OCI-Ly1, OCI-Ly7, SUDHL6) transduced with lentivirus expressing scrambled or short hairpin RNA (shRNA) against *IKZF1* (A), treated with increasing doses of lenalidomide (B), or combining shRNA with or without lenalidomide treatment (C). Proteins were measured by flow cytometry and normalized to scrambled transduced (A), DMSO-treated (B), or scrambled transduced and DMSO-treated (C) cells. RFI = relative fluorescence intensity. **(D)** IKAROS and CD19 median expression in isogenic IKAROS WT or KD CLL cell lines (CI, JVM-2, WA-OSEL). Proteins were measured by flow cytometry and normalized to WT condition. RFI = relative fluorescence intensity. **(E)** IKAROS and CD19 median expression in isogenic IKAROS WT or KD B-ALL cell lines (NALM6, REH, SUP-B15) used for ATAC-seq and RNA-seq experiments. Proteins were measured by flow cytometry and normalized to WT condition. RFI = relative fluorescence intensity. **(F)** Z-score of differentially expressed genes between isogenic IKAROS WT and KD B-ALL cells. **(G)** GSEA for Hay Bone Marrow Pro-B gene signature^69^ in IKAROS WT and KD B-ALL cells. Bar plots in (A - E) show mean ± SEM. Statistical tests used were two-way ANOVA followed by Tukey’s multiple comparisons test (A - C); and one-way ANOVA followed by Šidák’s multiple comparisons test (D - E). *P<0.05, **P<0.01, ***P<0.001, ****P< 0.0001.

**Supplementary Figure S5:**
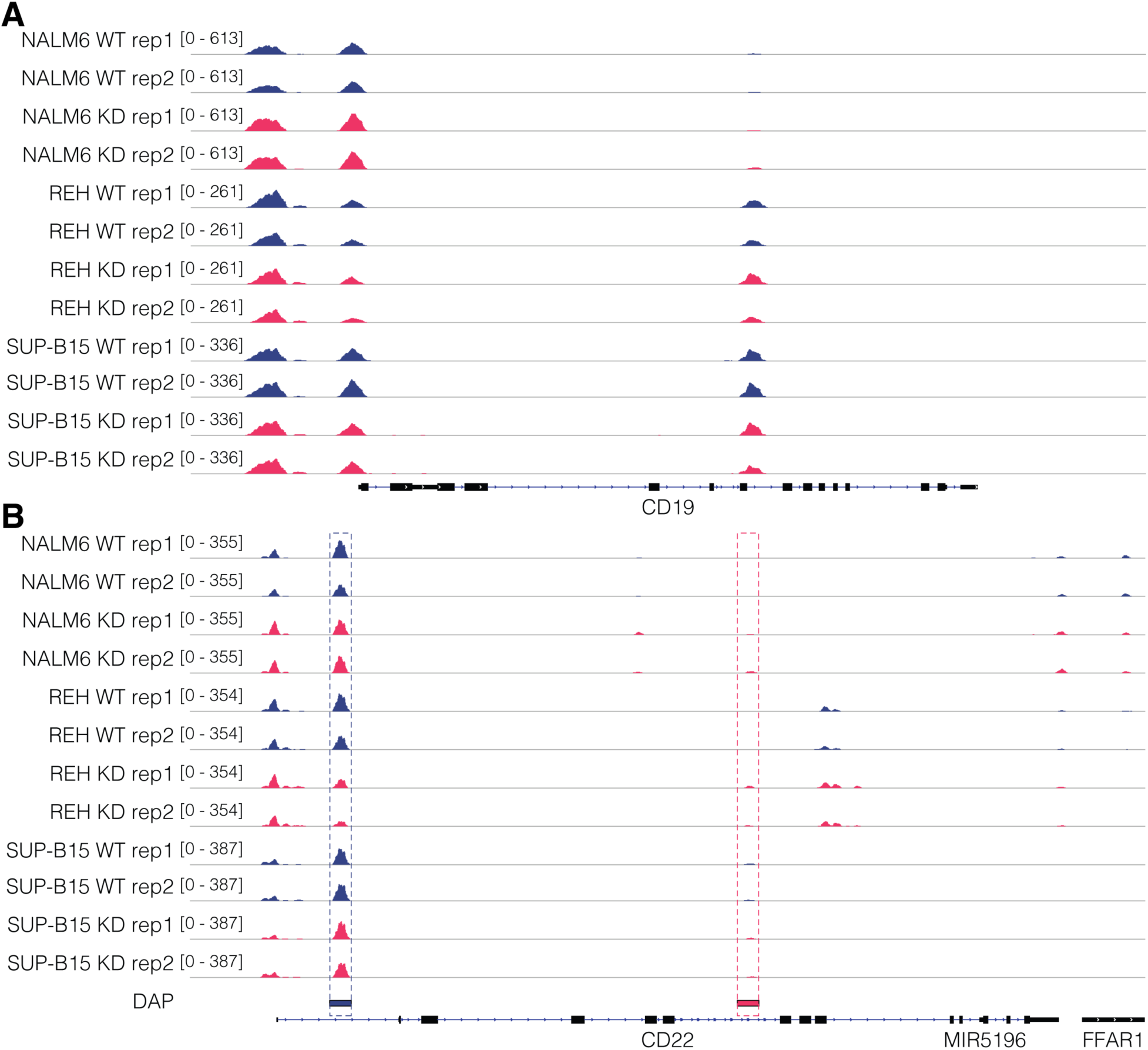
Low levels of IKAROS promote changes in chromatin accessibility at the CD22 gene, while not affecting the CD19 gene. **(A - B)** ATAC-seq chromatin accessibility track to *CD19* (A) and *CD22* (B) promoter and gene from isogenic IKAROS WT and KD B-ALL cell lines. Experiment was performed in 3 B-ALL cell lines (NALM6, REH, SUP-B15) in duplicate. DAP stands for differentially accessible peaks in IKAROS WT (blue) or KD (red) conditions.

**Supplementary Figure S6:**
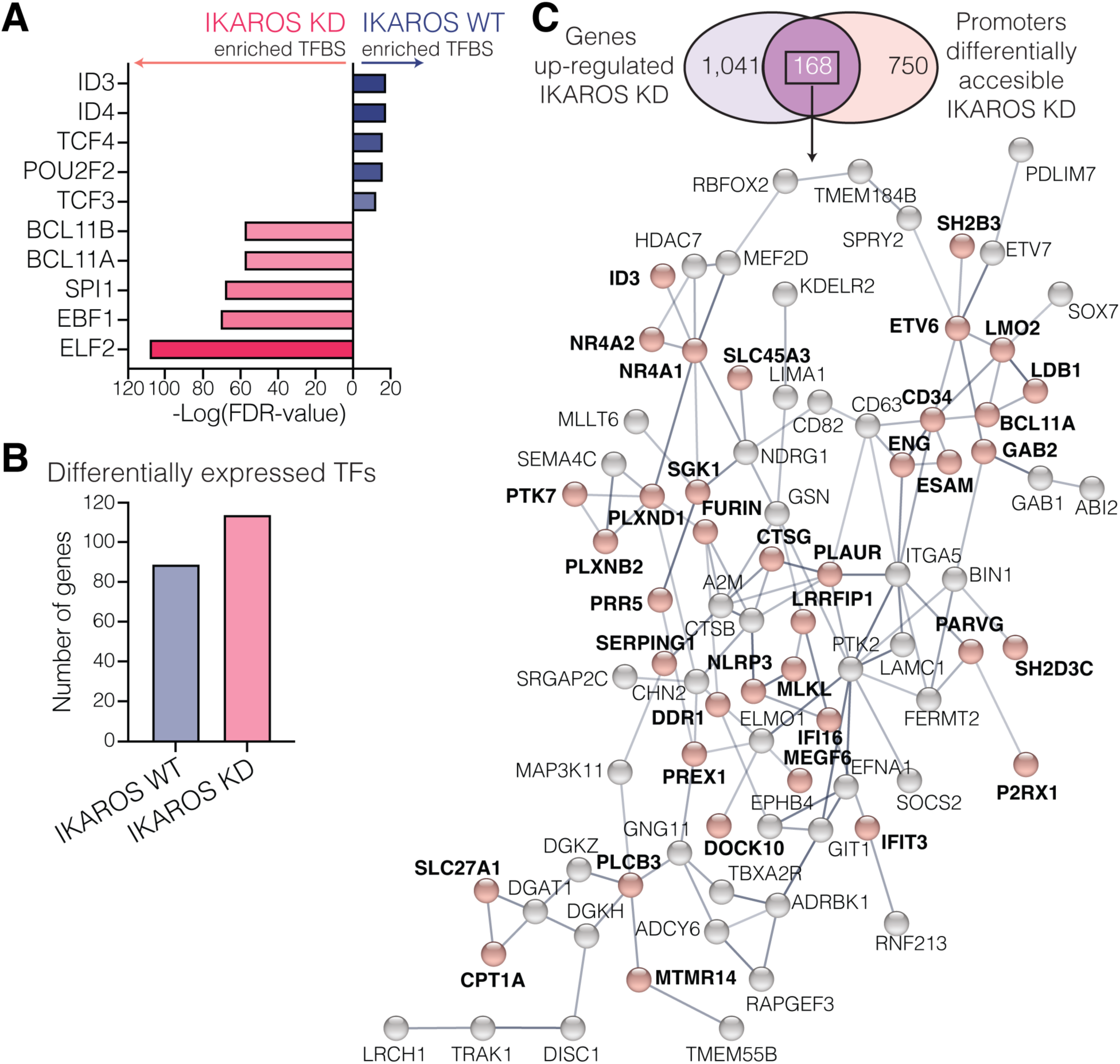
Low levels of IKAROS induces expression of a non-B lineage gene network. **(A)** Top 10 transcription factor binding sites (TFBS) enriched in differentially open peaks. **(B)** Number of differentially up-regulated transcription factor encoding genes in IKAROS WT or KD B-ALL cells. **(C)** Venn diagram between genes up-regulated and genes whose promoters have differentially accessible peaks in IKAROS KD B-ALL cells. STRING (Search Tool for Retrieval of Interacting Genes/Proteins) protein-protein interaction network of genes up-regulated and with more accessible promoters in IKAROS KD cells. Node represents gene and line thickness is proportional to the confidence of interaction. Red nodes correspond to genes associated with non-B cell lineage.

**Supplementary Figure S7:**
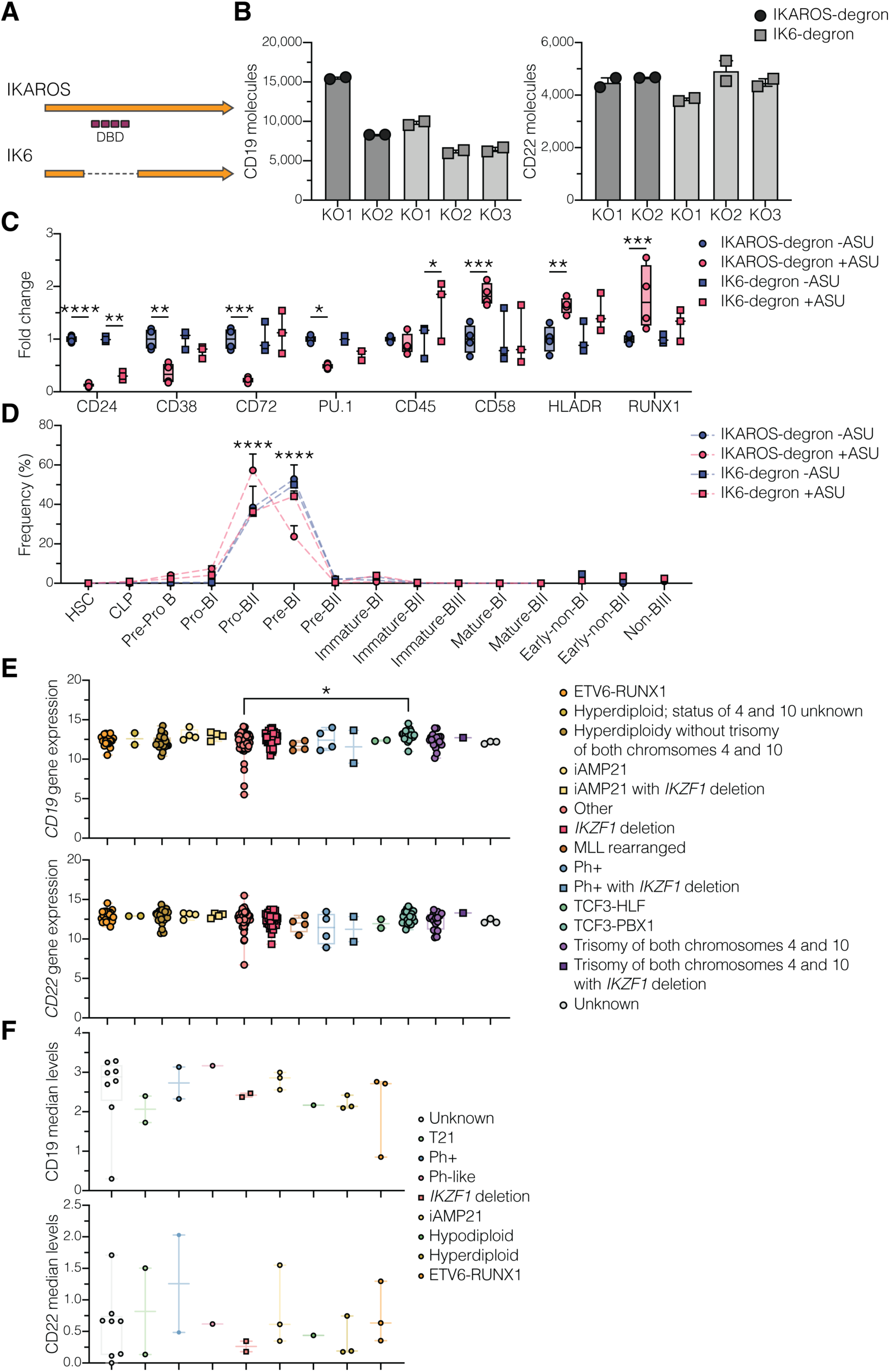
Deletion of IKAROS DNA binding domains abrogate modulation of CD19 and CD22 surface expression. **(A)** Schematic representation of IKAROS deletion used to generate the IK6-degron model. DBD stands for DNA binding domains. **(B)** Baseline CD19 (left) and CD22 (right) molecule numbers on the cell surface of IKAROS-degron (KO1 and KO2; circles) and IK6-degron (KO1, KO2, and KO3; squares) models. Experiment was performed in duplicate. **(C)** IKAROS- and IK6-degron models were treated with DMSO (-ASU; n = 4), or 10 µM asunaprevir (+ASU; n = 4) for 7 days. Protein profile was measured by CyTOF and normalized to -ASU condition. **(D)** Developmental classification of samples in (C). Frequencies of Pro-BII and Pre-BI populations were statistically different (p-value < 0.0001) in the IKAROS-degron +ASU condition compared to the other conditions. **(E)** *CD19* (top) and *CD22* (bottom) vst counts across different B-ALL genomic subtypes with or without *IKZF1* deletions in B-ALL patient samples (n = 282) from TARGET ALL project. **(F)** Bulk CD19 (top) and CD22 (bottom) median levels in pre-CART19 samples (Figure 1A) based on their cytogenetic classification. Bar plots in (B) show mean ± SEM. Boxplots in (C), and (E – F) show median with range from minimum to maximum values. Curves in (D) show mean ± SEM. Statistical tests used were Kruskal-Wallis followed by Dunn’s multiple comparisons tests (B) and (F); two-way ANOVA followed by Tukey’s multiple comparisons tests (C - D); and one-way ANOVA followed by Tukey’s multiple comparisons tests (E); *P<0.05, **P<0.01, ***P<0.001, ****P< 0.0001.

**Supplementary Figure S8:**
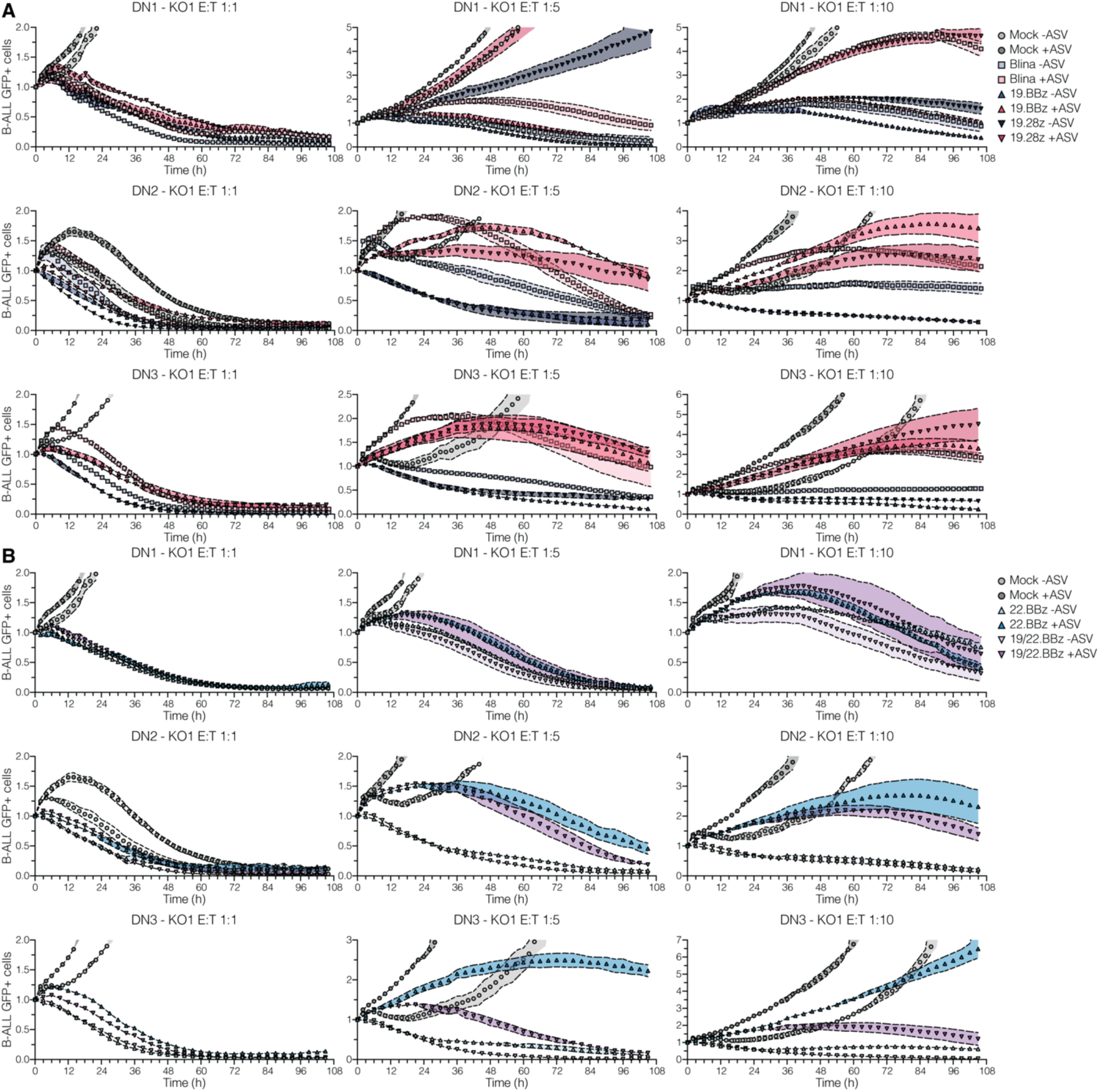
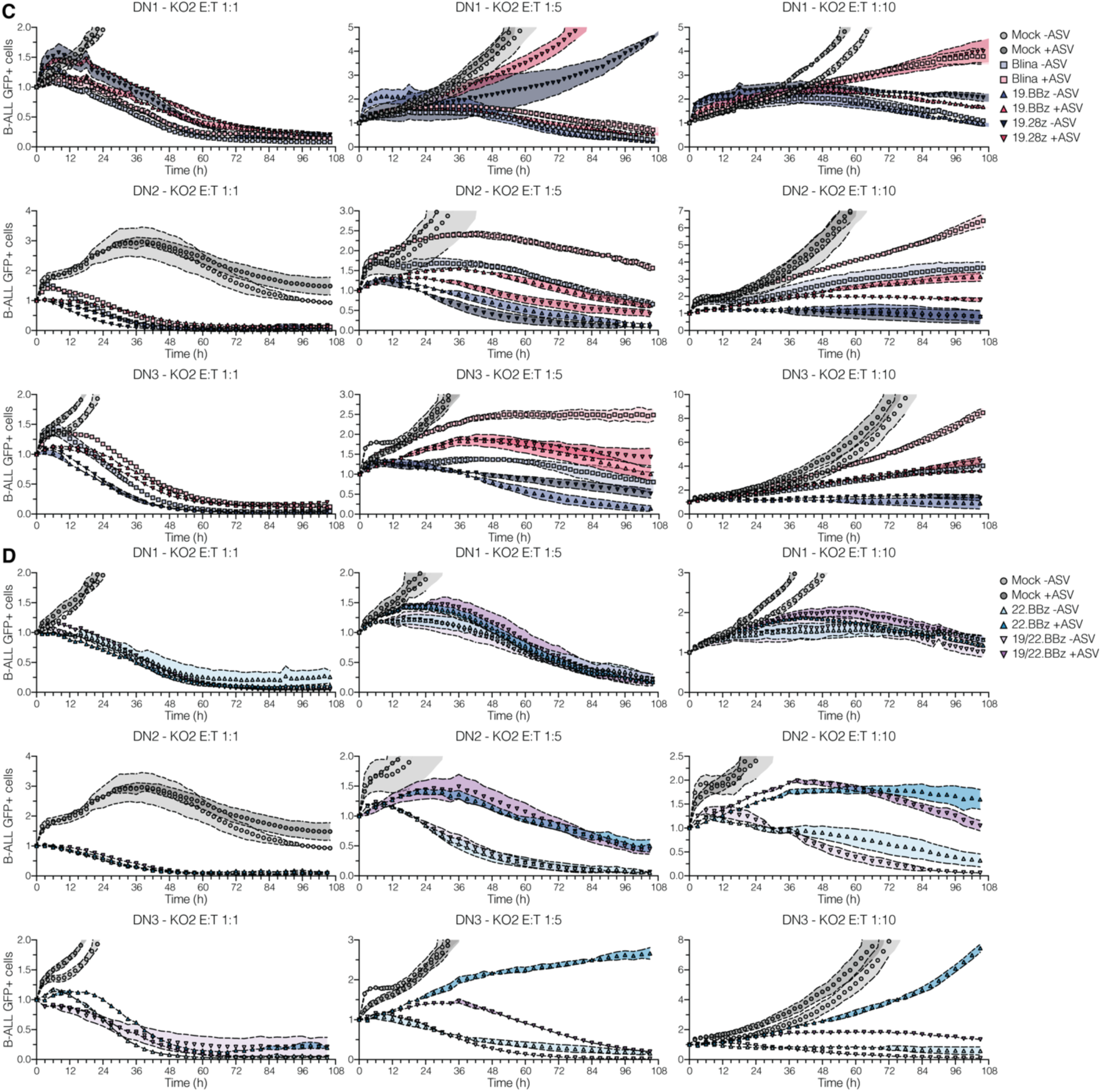
Low IKAROS levels confer resistance against CD19- and CD22-targeted therapies. **(A - D)** IKAROS-degron models (KO1, A - B; and KO2, C - D) were treated with DMSO (-ASU) or 10 µM asunaprevir (+ASU) for 7 days, before being co-cultured with mock, blinatumomab-treated, 19.BBz, and 19.28z (A or C) CAR T cells; or mock, 22.BBz, and dual 19/22.BBz (B or D) CAR T cells at 1:1, 1:5, and 1:10 E:T ratio. Experiment was conducted in triplicate, with different T cell donors for each replicate. B-ALL cell viability was measured at every 2 - 3 h interval via IncuCyte. GFP median values were normalized to 0 h condition. RFI = relative fluorescence intensity. Curves in (A - D) show mean ± SEM. Dots represent the mean value from three technical replicates.

**Supplementary Figure S9:**
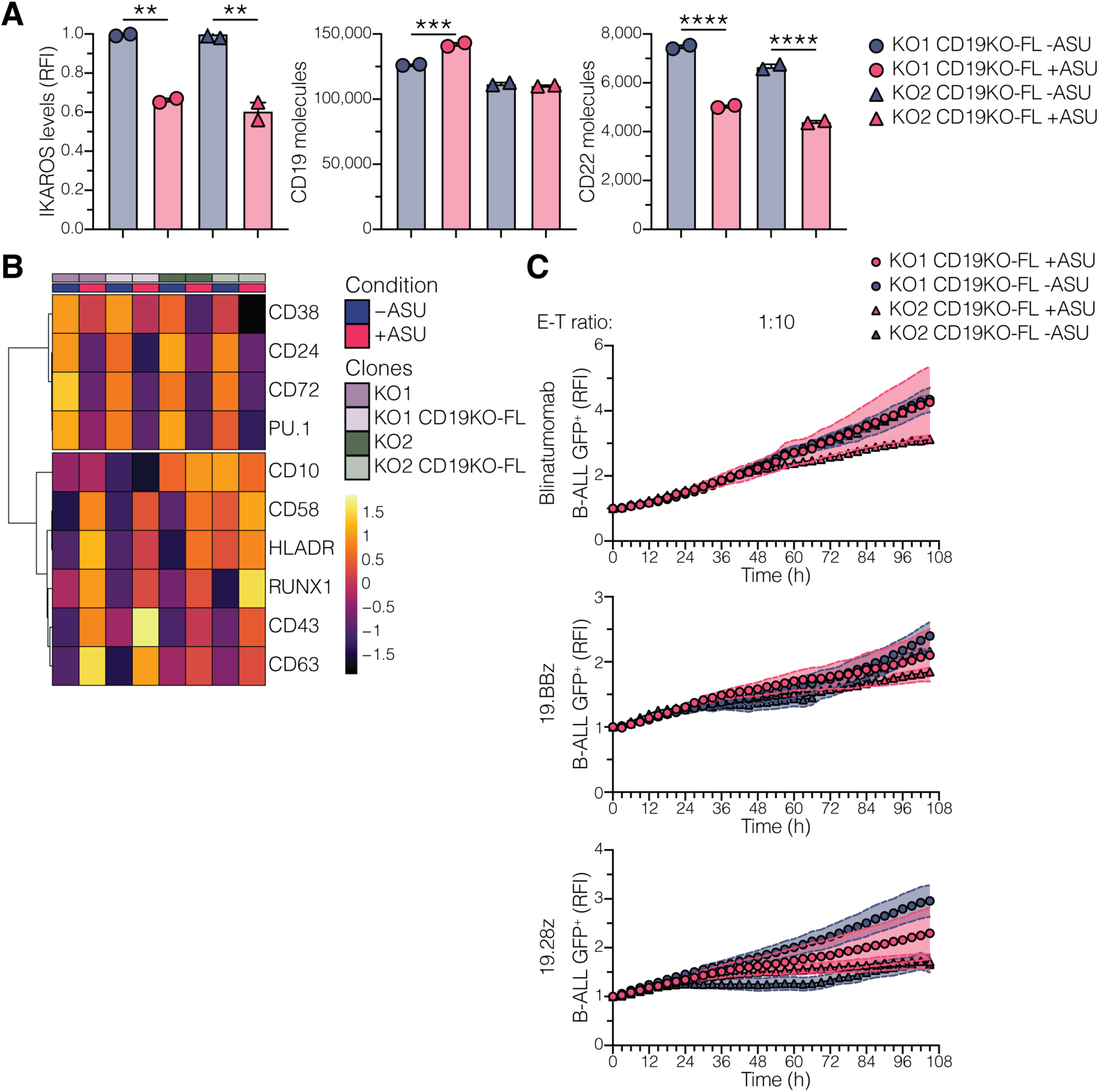
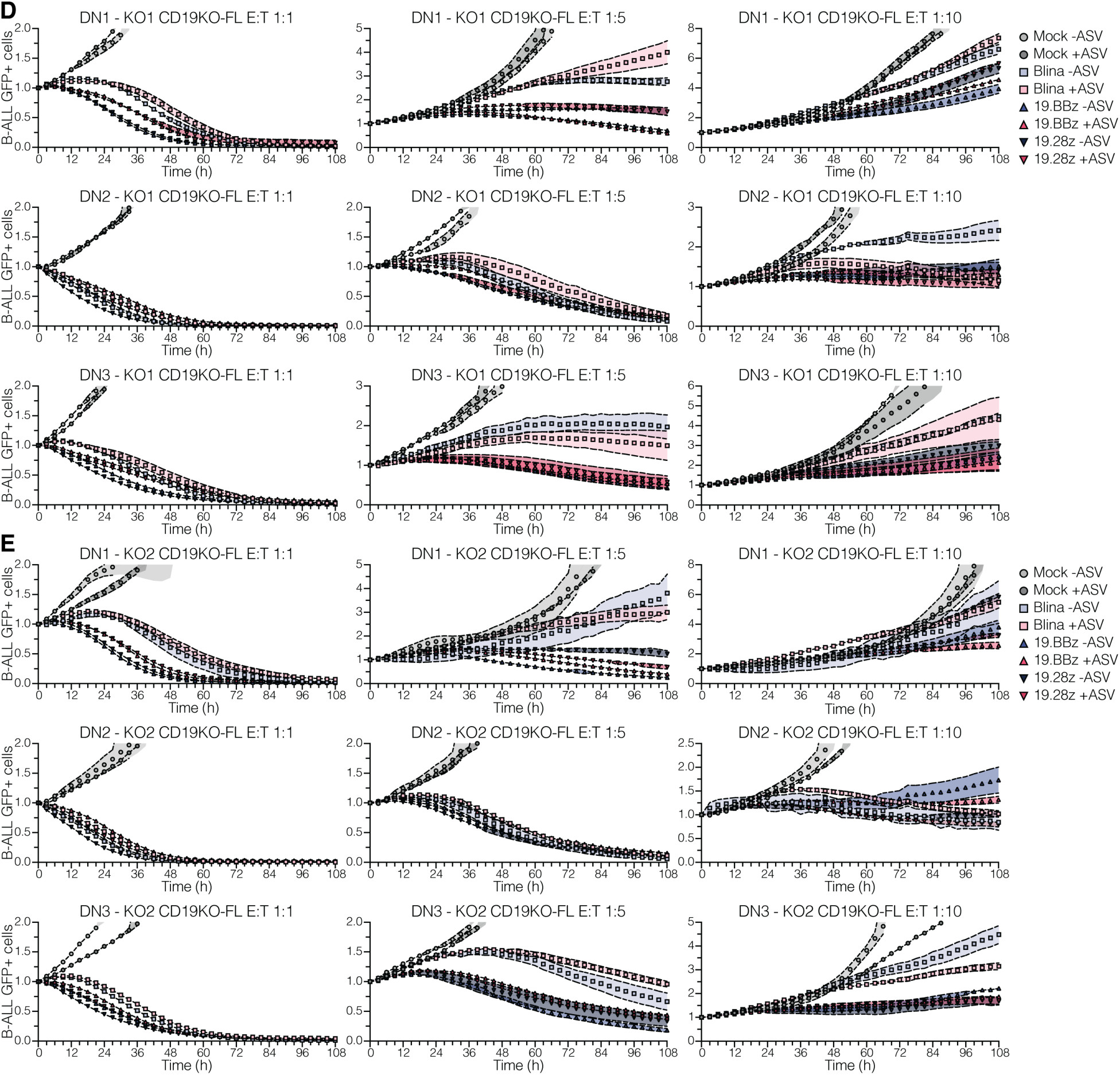
IKAROS depends on CD19 modulation to confer resistance to CD19-targeted therapies. **(A)** Relative IKAROS median levels (left), CD19 (middle), and CD22 (right) molecule numbers in IKAROS-degron CD19KO-FL model treated with DMSO (-ASU) or 10 µM asunaprevir (+ASU) for 7 days. Values were measured by flow cytometry. IKAROS median values were normalized to -ASU condition. RFI = relative fluorescence intensity. **(B)** Median expression of key proteins in IKAROS-degron and IKAROS-degron CD19KO-FL models treated with DMSO (-ASU) or 10 µM asunaprevir (+ASU) for 7 days. **(C)** IKAROS-degron CD19KO-FL models treated with DMSO (-ASU) or 10 µM asunaprevir (+ASU) for 7 days, and co-cultured with blinatumomab-treated (up), 19.BBz (middle), and 19.28z (bottom) CAR T cells at 1:10 E:T ratio. **(D - E)** IKAROS-degron CD19KO-FL models (KO1, D; and KO2, E) treated with DMSO (-ASU) or 10 µM asunaprevir (+ASU) for 7 days, and co-cultured with mock, blinatumomab-treated, 19.BBz, and 19.28z CAR T cells at 1:1, 1:5, and 1:10 E:T ratio. Experiment was conducted in triplicate, with different T cell donors for each replicate. B-ALL cell viability was measured at every 2 - 3 h interval via IncuCyte. GFP median values were normalized to 0 h condition. RFI = relative fluorescence intensity. Bar plots in (A) show mean ± SEM. Curves in (C - E) show mean ± SEM. Dots represent the mean value from three technical replicates. Statistical tests used were one-way ANOVA followed by Tukey’s multiple comparisons tests (A); and two-way ANOVA followed by Šidák’s multiple comparisons test (C - E). **P<0.01, ***P<0.001, ****P< 0.0001.

